# A chromatin fiber model explains cell-free DNA fragmentation signatures of active regulatory elements

**DOI:** 10.1101/2025.11.06.686988

**Authors:** Alexis Yang, Garyoung Gary Lee, Surya Chhetri, Razane El Hajj Chehade, Gunsagar Gulati, Medha Pandey, Doris Fu, Yoo-Na Kim, Shahab Sotudian, Hunter Savignano, Sadia D. Shahbazi, Chelsea Philpot, Taylor A. Hoggood, Doga C. Gulhan, Manolis Kellis, Jacob E. Berchuck, Sylvan C. Baca

## Abstract

Circulating cell-free DNA (cfDNA) assays are being widely adopted in oncology and maternal-fetal medicine. Patterns of cfDNA fragmentation can provide useful information about gene regulation and expression in human disease from a blood draw. Here, we demonstrate that enhancer RNA expression – a marker of enhancer activity – can be inferred from local patterns of cfDNA fragmentation. We define a transcriptional activation score (TAS) that predicts expression of enhancers and genes based on cfDNA fragment sizes and positions near transcriptional start sites (TSSs). The TAS identifies activity of cancer-associated enhancers in patients with cancer, distinguishes clinically relevant cancer subtypes, and identifies activation of enhancers associated with treatment resistance and therapy response. We propose a simple model to account for our findings based on chromatin fiber structure and the depletion of H1 histone proteins near active TSSs. Our model provides a unified framework that reconciles seemingly conflicting observations from prior fragmentomics studies. Broadly, this work enables blood-based assessments of gene regulation in cancer and non-oncologic diseases to inform pathobiology, diagnosis, and treatment selection.

## Introduction

Circulating cell-free DNA (cfDNA) tests are being adopted into diagnostic workflows for maternal-fetal medicine and oncology. Sequencing cfDNA can provide actionable molecular information in pregnancy and cancer from a simple blood draw^1,2^. The first generation of cfDNA assays focused on detecting genomic alterations in cancer and chromosomal abnormalities in pregnancy, but recent methods are extending the reach of cfDNA by providing information about gene regulation.

Recently, a second generation of cfDNA assays has emerged that focus on patterns of cfDNA fragmentation – collectively termed “fragmentomics”. Fragmentomics studies have identified cfDNA fragment features associated with the presence of cancer, including biases toward shorter and more variable fragment lengths and enrichment of nucleotide motifs at fragment ends^3–7^. These “global” differences in cfDNA fragmentation patterns in cancer may be due activation of specific nucleases in cancer cells during apoptosis or cell death from necrosis^5,8^. Importantly, cfDNA fragmentation is also influenced by “local” chromatin structure, reflecting the presence of histone modifications, open chromatin, expressed gene promoters, intron-exon boundaries at expressed genes, and/or bound transcription factors (TFs)^9–18^.

How gene regulation and local chromatin structure influence cfDNA fragmentation is a central question in the study of cfDNA. Answering this question could enable accurate non-invasive inference of gene regulation and expression in disease, considerably extending the clinical utility of cfDNA assays. Several studies have linked fragment length patterns to gene expression or regulation, but the underlying mechanisms in some cases are unclear, and some findings appear contradictory. For example, studies have reported enrichment of both long (*e.g.*, > 200 bp)^19^ and short fragments (< 100bp) at active promoters, as well as increased fragment size variability^14^. A recent study reported shorter fragments in bodies of highly expressed genes^20^, while another associated shorter cfDNA fragments with inaccessible chromatin^21^. Further demonstrating the complexity of these associations, a recent paper revealed intricate biases in sizes of cfDNA fragments at active regulatory elements marked by the histone modification H3K27ac^12^. Overall, the mechanistic basis for these observations remains unclear, as studies have yet to examine the full range of cfDNA fragment features associated with regulatory element activity and explain their origins in a unified framework.

We set out to systematically examine the associations between cfDNA fragmentation patterns and gene regulation by focusing on enhancers. Enhancers are promoter-distal regulatory elements that are bound by TFs and form loops with gene promoters to regulate transcriptional activity^22–24^. Enhancers control context-dependent gene expression programs that are essential for organismal development, cellular differentiation, and response to environmental stimuli^23–26^. By capturing epigenetic cellular “states” that are shaped by gene regulation, enhancer profiling can provide insights into the pathogenesis of human disease.

Upon activation, enhancers are transcribed, producing enhancer RNAs (eRNAs) – short, non-coding transcripts that are required for enhancer function and serve as dynamic markers of enhancer activity^25,27,28^. eRNAs are quickly degraded and, due to their short half-life, provide real-time information about enhancer activation status^27,29,30^. While recent advances in fragmentomics have demonstrated the ability to infer gene expression from cfDNA fragmentation patterns^13,14^, whether these features can delineate enhancer expression and activity is unknown.

Here, we show that enhancer expression can be inferred from cfDNA fragmentation patterns. We identify complex associations of cfDNA fragment size and locations near transcriptional start sites (TSSs) that correlate tightly with expression of both eRNAs and genes. With generative modeling of cfDNA cleavage, we identify a mechanistic basis for these fragmentation patterns, which are explained by nucleosome repeat length and the presence of H1 linker histone protein, two determinants of chromatin fiber structure. This model explains several observations from prior fragmentomics studies and enables accurate inference of expression for clinically relevant genes and enhancers. Our findings establish a versatile platform for measuring gene regulation from cfDNA.

## Results

### Dinucleosome cfDNA fragments are enriched at active enhancers

To investigate how cfDNA fragmentation reflects enhancer activity, we analyzed cfDNA fragments from H3K27ac cell-free chromatin immunoprecipitation (cfChIP-seq) experiments, which enrich nucleosomes from active regulatory elements. We analyzed the fragment length distributions from H3K27ac cfChIP-seq data from 7 healthy individuals and compared them to fragment length distributions in cfDNA whole genome sequencing (WGS) data from the same samples^31^. cfDNA WGS fragment sizes showed the expected mode near 167 bp, corresponding to the length of DNA wrapped around histones to form a mononucleosome, and a small second peak near 320 bp, corresponding to dinucleosome-sized fragments (**Fig. 1a**). In contrast, cfDNA fragments enriched by H3K27ac cfChIP-seq showed a marked elevation in the portion of dinucleosome fragments and a decrease in mononucleosome fragments (**Fig. 1a**). We observed a similar, though less pronounced, increase in the dinucleosome fragments and decrease in mononucleosome fragments in cfDNA WGS data at H3K27ac-marked regions, indicating that the differences in dinucleosome and mononucleosome representation was not solely attributable to immunoprecipitation (**Fig. 1a**). In addition, the modal dinucleosome fragment sizes were shorter in H3K27ac cfChIP-seq data and cfDNA WGS data at H3K27-acetylated regions (median sizes 311 and 314 bp) compared to unselected WGS fragments (median size 322 bp; p-value = 8.7 x 10^-4^, and 3.0 x 10^-3^, respectively; one-sided Wilcoxon rank sum test; **Fig. 1b**). These results indicate that dinucleosome cfDNA fragments from active chromatin are shorter and more abundant than dinucleosome fragments from the rest of the genome, which we investigated further.

**Figure 1.**
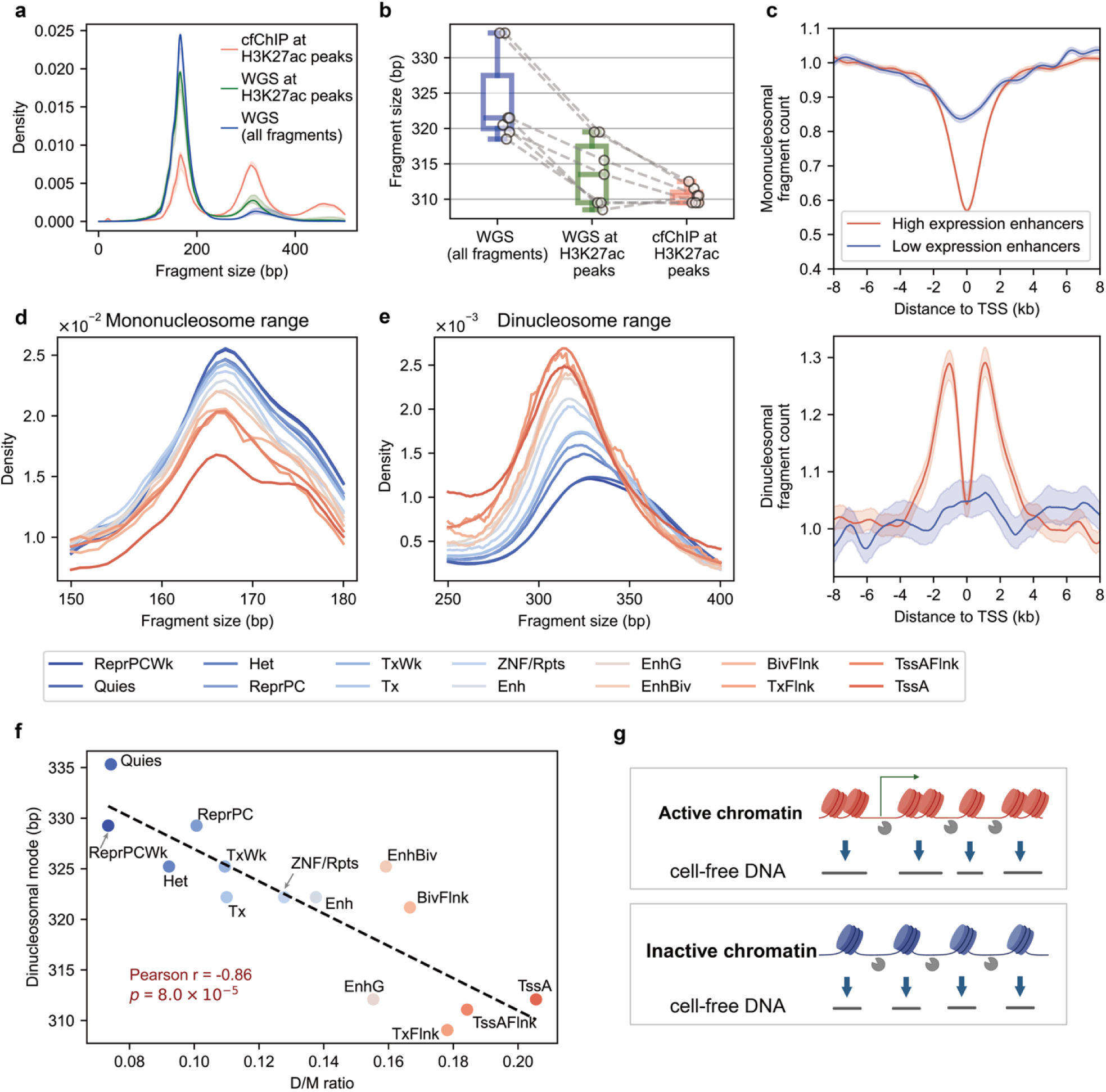
Tighter nucleosome spacing increase the ratio of dinucleosome to mononucleosome sized fragments near expressed enhancers. **(a)** Distributions of cfDNA fragment lengths from N=7 healthy volunteers. H3K27ac cfChIP-seq fragments are shown in orange and compared to WGS fragments at H3K27ac peaks identified from cfChIP-seq data (green) and to unselected WGS fragments (blue) from the same plasma samples. Shaded regions represent the interquartile range (IQR) across samples. **(b)** Comparison of the modal dinucleosome size, defined as the local maximum of the peak near 300bp in **(a)**. Each dot represents one sample. Gray dashed lines connect observations from the same samples. **(c)** Aggregated midpoint count of mononucleosome size (140-180 bp; upper panel) and dinucleosome size (300-340 bp; lower panel) fragments near enhancer transcriptional start sites (TSS). Shaded regions represent 95% bootstrap confidence intervals. **(d)** Mononucleosome size fragment distributions from pooled healthy plasma sample for various chromatin status. **(e)** As in **(d)**, but dinucleosome size fragment distribution. For visualization purposes, values were smoothed by averaging across bins of bp (d) and 10bp (e). **(f)** Fragment size vs. the ratio of dinucleosome size to mononucleosome size fragments (D/M ratio) for cfDNA mapping to indicated chromHMM states^35^ (Methods). **(g)** A model accounting for increased D/M ratio and shorter dinucleosome fragment size in cfDNA fragments from active vs. inactive chromatin.

We next asked whether the differences in dinucleosome fragment size and abundance are present in cfDNA from enhancers with high eRNA expression, which tend to be marked by H3K27ac^32^. We analyzed dinucleosome and mononucleosome coverage in cfDNA WGS data near enhancer TSSs with varying levels of eRNA expression, as measured by PRO-cap in lymphoblastoid cell lines^33^. Because hematopoietic cells are the main source of cfDNA in healthy individuals, these cells provide a relevant proxy for assessing enhancer-associated fragmentation patterns. Near the TSS of highly expressed enhancers, dinucleosome coverage increased markedly, while mononucleosome coverage decreased (**Fig. 1c**). A similar pattern was observed at promoter TSSs (**Fig. S1**). These results indicate a focal cfDNA fragmentation pattern at transcribed regulatory elements, with increased dinucleosome fragments flanking the TSS and decreased mononucleosome fragments at the TSS.

Prior studies of nucleosome positioning in cell lines have shown that chromatin with active histone modifications (*e.g.*, H3K27ac) is associated with tight nucleosome spacing, while regions marked by repressive modifications (*e.g.*, H3K27me1) tend to exhibit looser nucleosome spacing^34^. We hypothesized that such differences in nucleosome spacing could account for the increase in dinucleosome fragments observed at transcriptionally active sites. To test this, we defined a set of consensus chromatin states in blood cell types. We used ChromHMM chromatin state annotations from 27 hematologic cell types from the Roadmap Epigenomics Project^24,35^ and focused on sites where more than half of all samples shared the same chromatin state. We analyzed mono- and di-nucleosome proportions of pooled 7 healthy individuals across chromatin states (**Fig. 1d** and **e**). A striking difference in mono- and di-nucleosome proportions was observed across chromatin states. cfDNA from active chromatin had more dinucleosome-sized fragments and fewer mononucleosome-sized fragments. Across chromatin states, we observed a strong negative correlation between the ratio of dinucleosome to mononucleosome-sized fragments (D/M ratio) and the modal dinucleosome sizes (Pearson’s r = −0.86; p = 8.0x10^-5^; **Fig. 1f**). For example, active states such as intragenic enhancers (EnhG) and active transcriptional start sites (TssA) had shorter dinucleosome sizes and larger D/M ratios (305-315 bp, D/M ratio: >0.15), while quiescent and repressive states exhibited larger dinucleosome sizes and smaller D/M ratios (325-335 bp, D/M ratio: <0.1). These findings indicate that dinucleosome cfDNA fragments originating from more transcriptionally active or accessible chromatin are both shorter in size and more abundant. Further, they point to a genome-wide inverse relationship between D/M ratios and dinucleosome fragment size, which we sought to explain.

We considered mechanistic explanations for the shorter dinucleosome sizes and increased D/M ratios in cfDNA from active regulatory elements. One possible explanation is that nucleosomes in active chromatin are more tightly spaced, leaving less exposed linker DNA between adjacent nucleosomes (**Fig. 1g**). In this model, reduced exposure of linker DNA decreases the chance of inter-nucleosomal cleavage, preserving more dinucleosome-sized fragments. These dinucleosome fragments are shortened due to smaller distances between nucleosomes. In contrast, inactive regions exhibit wider nucleosome spacing, which exposes more inter-nucleosomal linker DNA, increasing the likelihood of cleavage between nucleosomes and leading to a greater proportion of mononucleosome-sized fragments. This model aligns with our observation that increased enhancer activity is associated with shorter, more abundant dinucleosome-sized fragments.

### cfDNA fragment sizes and locations correlate tightly with transcription

Intrigued by the focal elevation of D/M ratios near expressed enhancers, we systematically investigated how cfDNA fragment sizes correlate with enhancer expression. We computed the correlation between the abundance of fragments grouped by length and eRNA expression across a range of distances from enhancer TSSs. Using pooled healthy LP-WGS cfDNA data^31^ (∼15x coverage from 34 samples), we observed highly localized, distance-dependent relationships between fragment size and transcript levels (Fig. 2a). At the enhancer TSS, sub-nucleosomal (<120 bp) and intermediate-size (∼200–300 bp) fragments were positively correlated with eRNA expression (r = 0.89 and 0.86, respectively), while mononucleosome fragments (∼170 bp) exhibited a strong negative correlation (r = –0.83). These patterns changed in the enhancer “shoulder” regions (>1 kb from the TSS), where dinucleosome-sized fragments (∼310 bp) were enriched in active enhancers, while mononucleosomes remained depleted. In all, 55 % of fragment size–distance combinations within 2kb of enhancer TSSs were significantly correlated or anti-correlated with expression at a false-discovery rate of 0.05). These findings reveal relationships between fragment size and eRNA expression that are highly position-dependent, with distinct fragmentation patterns at the TSS compared to the shoulder regions.

**Figure 2.**
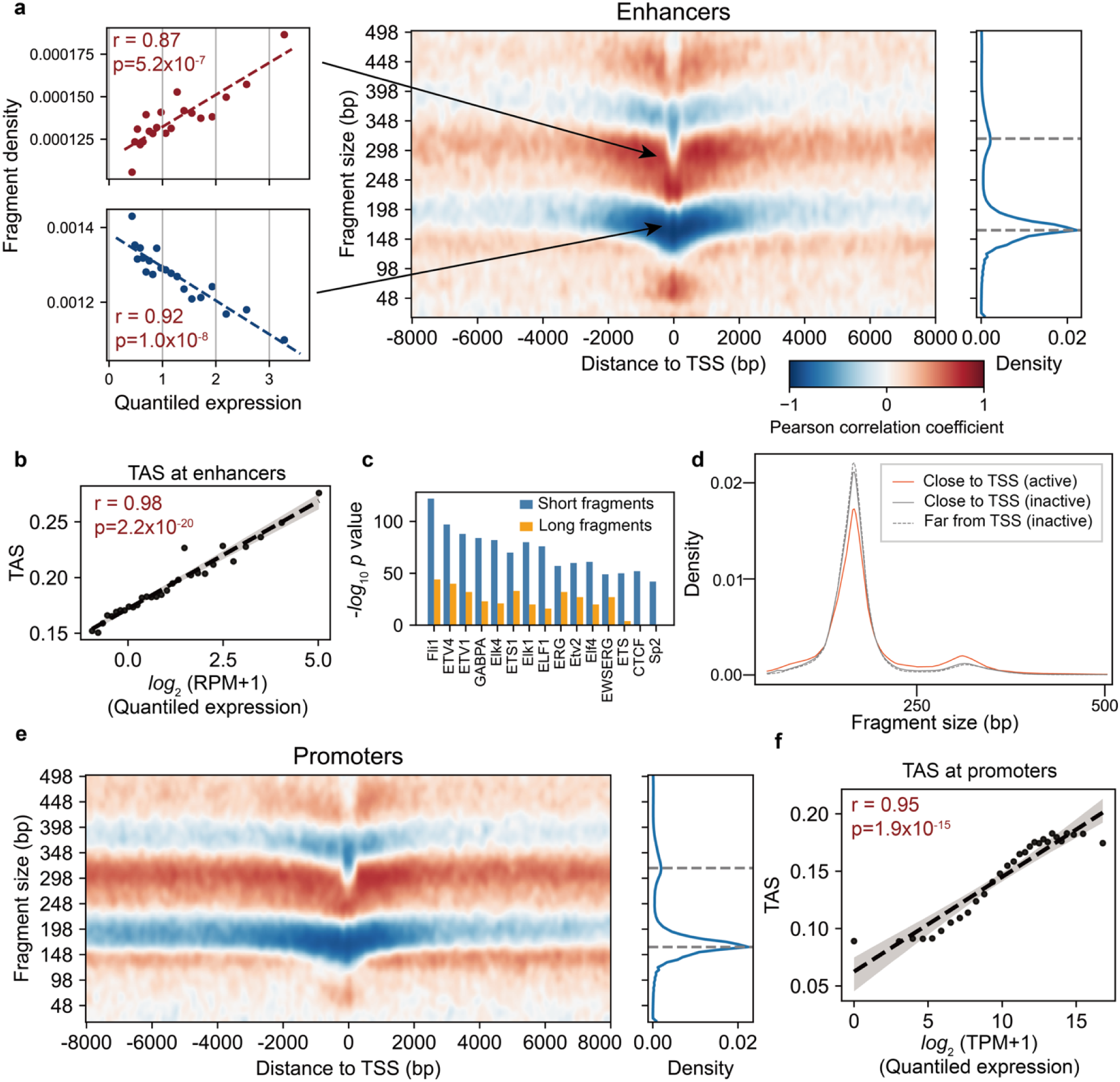
Consistent patterns of correlation between cfDNA fragmentation and transcription at promoters and enhancers. **(a)** Healthy volunteer samples (N=34) were aggregated (N=286M fragments, in total) and grouped by fragment size and distance to TSS. The number of fragments in each group was normalized to the total number of fragments within 8kb of enhancer TSSs for each sample and tested for correlation with enhancer expression. Enhancers and are ranked by expression and aggregated into N=30 groups. Scatterplots (left) show examples for modal fragment size categories: mononucleosomal (∼150-170 bp, red) and dinucleosomal (∼300-320 bp, blue) fragments within +/-1 kb of enhancer TSSs. Correlation values are plotted in the heatmap (middle). The right figure shows the fragment size distribution of aggregated sample at enhancer regions. Dashed gray lines refers mononucleosome and dinucleosome fragment sizes. **(b)** Correlation between TAS values at enhancers and eRNA expression values. Enhancers (N=37,094) were aggregated along with their eRNA expression quantile. TAS of pooled LP-WGS data (N=32) were calculated for aggregated enhancers. The RPM stands for read per million. **(c)** Top TF motifs enriched within long (147-500 bp) vs. short (<120 bp) fragments within 500 bp from the TSS were presented with their enrichment *p*-values. **(d)** Distributions of active and inactive cfDNA fragment sizes at sites close to (<500 bp) and far from (2000-3000 bp) the enhancer TSS. **(e)** and **(f)** As in (a) and (b) but showing TSSs and expression correlation values for N= 17,424 genes. The TPM stands for transcript per million.

Based on these observations, we developed a transcriptional activation score (TAS) that integrates fragment size and positional correlation patterns. The TAS is computed as a weighted ratio of positively and negatively correlated fragment bins and serves as a compact index of enhancer or promoter activity (Methods, **Fig. S2, Supplementary tables 1 and 2**). Applying this metric to an independent cohort of pooled lung cancer LP-WGS samples^36^ with 0% estimated tumor fraction by ichorCNA^37^ (n = 32, total pooled coverage = 16.3x), we found that TAS was strongly associated with eRNA expression (Pearson r = 0.98; **Fig. 2b**). This finding demonstrates the generalizability of the correlations we identified between fragmentation and expression as well as the robustness of the TAS.

We considered gene regulatory mechanisms that could underlie the correlation patterns identified by the TAS. We hypothesized that the enrichment of <100 bp fragments near highly active enhancers likely reflects nucleosome depletion and protection by bound transcription factors (TFs)^11,17^. To test this, we examined motif enrichment in short (<100 bp) versus longer fragments near the TSSs of enhancers highly expressed in lymphoblastoid cells. Short fragments showed significantly stronger enrichment of canonical TF motifs, including motifs corresponding to the megakaryocyte lineage TF FLI1^38^ and the lymphoid-associated TF ETS1^39^ (**Fig. 2c**), supporting a model in which these short cfDNA fragments are protected from nucleases by bound TFs^11,17^. In addition, highly expressed enhancers had more diverse fragment lengths at their TSSs compared to shoulder regions or inactive chromatin (**Fig. 2d**), with fewer mono-nucleosome-size fragments and an increase in intermediate-length fragments (i.e., 225-300 bp). This finding accounts for high correlation between expression and the density of fragment between mono- and dinucleosome sizes, and is consistent with increased “fragmentation entropy” due to nucleosome repositioning at the active TSSs^14^.

Comparing fragmentation correlation patterns at enhancers with those at promoters, we observed similar behavior near promoter TSSs (**Fig. 2e**), and strong correlation between mRNA expression and TAS (Pearson r = 0.95; **Fig. 2f**), but with notable differences. First, the correlation of shorter TF-bound fragments (<100bp) was greater at enhancer TSSs, consistent with the observation that TFs often bind preferentially to enhancers^40,41^. Second, correlation patterns were less symmetric with respect to the TSSs for promoters (**Fig. S3**), which we speculate reflects unidirectional transcription of genes compared to bidirectional transcription of enhancers. In addition, correlation patterns decayed more rapidly with distance from the TSS for enhancers compared to promoters. Promoters maintained a strong positive correlation between dinucleosome fragments and expression beyond 8 kb from the TSS, while this correlation decayed at enhancer beyond ∼2 kb, corresponding to the maximum length of most enhancer RNAs^23,25^. This difference may reflect the short transcript length of enhancers compared to genes and suggests that the correlation patterns we observe associate with the genomic footprint of transcription.

### A chromatin fiber model accounts for cfDNA fragmentation patterns at active enhancers

We were intrigued by the correlation patterns on the “shoulders” of promoter and enhancer TSSs (e.g. > 500bp from the TSS), which differed strikingly from patterns at the TSS. We hypothesized that the association at the TSS shoulders between fragment size and expression could be explained by underlying structural differences between active and inactive chromatin. Two major determinants of chromatin fiber structure are the nucleosome repeat length (NRL) and the presence or absence of the H1 linker histone protein. The NRL has been shown to differ between transcriptionally active and silent regions in both structural studies of chromatin fibers and MNase-seq experiments^34,42–44^. H1 linker protein is depleted near active TSSs and can affect the structure of chromatin fibers, as well as the extent of cfDNA protected from degradation by a nucleosome^17,44–46^.

We built a biologically motivated generative model to ask whether simple differences in nucleosome spacing and histone-DNA interaction can explain the complex correlations of cfDNA fragment size and transcription. Briefly, the model simulates cleavage of DNA that is not protected by contact with a histone protein (Methods). The model tests a grid of biologically plausible values for two parameters: effective nucleosome size (the amount of DNA protected by a histone) and NRL (**Fig. 3a**). This model assumes normally distributed effective nucleosome sizes and NRLs, with different mean values for these parameters for active and inactive chromatin. For each parameter set, the relative abundance of fragments of a given size is compared between active and inactive chromatin simulations and plotted as a log ratio (**Fig. 3b**). We then compare these log ratios (“simulation”) to the correlation of a given fragment size with expression near promoters (“observation”) and identify parameters that maximize the correlation between simulation and observation (**Fig. 3b**).

**Figure 3.**
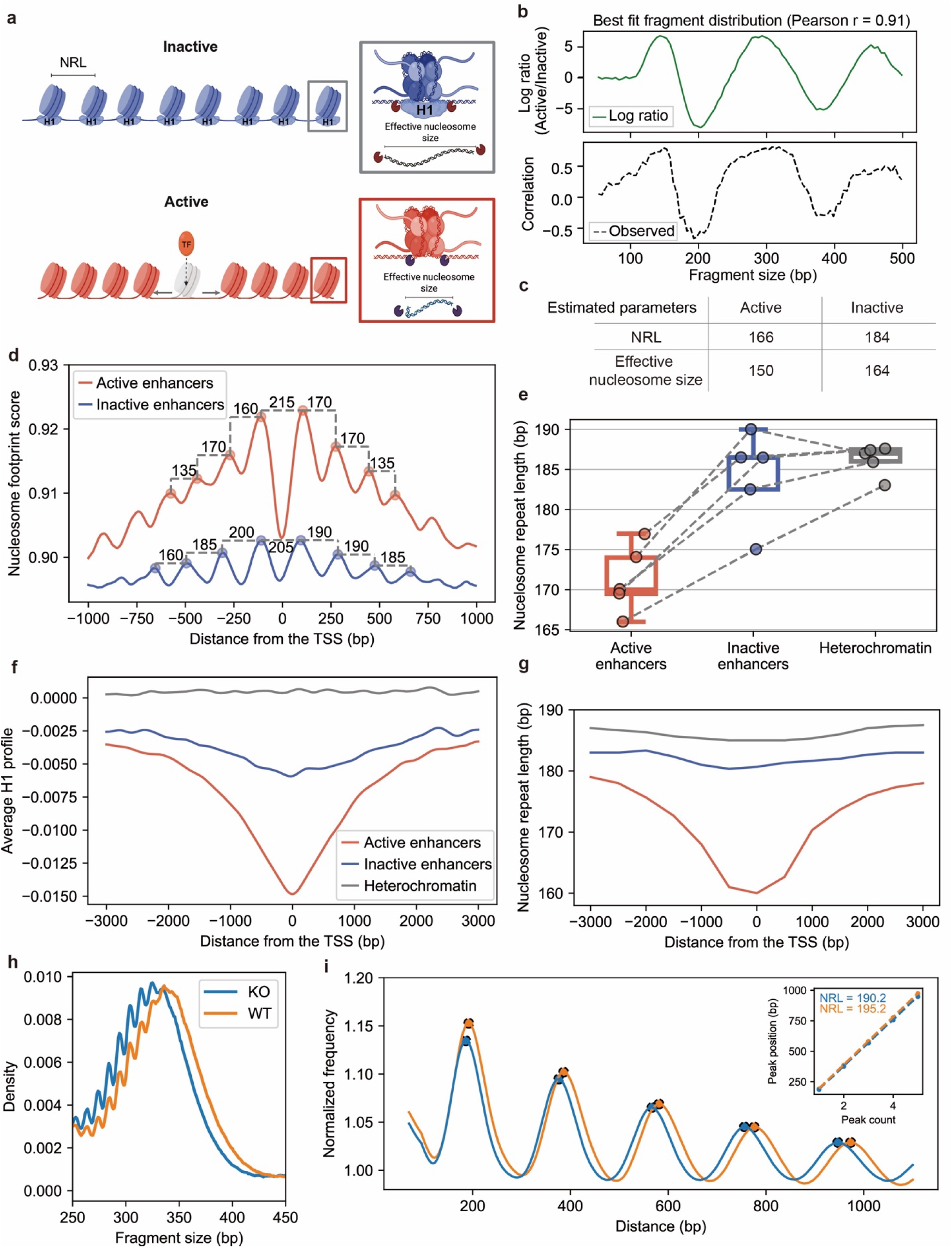
Integrated model of DNA fragmentation near active transcriptional start sites. **(a)** Parameters for the generative model of cfDNA fragmentation: nucleosome repeat length (NRL) and effective nucleosome size. **(b)** Correlation of model predictions with observed data. Dashed line indicates the observed correlation between expression and the number of fragments of the indicated size at +/- 3-5kb from a promoter TSS. Green line indicates the log-ratio of fragment counts of each size at active vs. inactive chromatin in the simulation. See methods for additional details. **(c)** The parameters for NRL and effective nucleosome size that maximize correlation between the observed data and simulations. **(d)** Nucleosome footprint score (NFS) for a single cancer-free plasma sample, and high-coverage cfDNA sequencing data (81x). NFSs were calculated across 5,000 aggregated enhancers with high expression and 5000 enhancers with zero expression in lymphoblastoid cell lines^33^. **(e)** Comparison of NRL near the enhancers with high vs. zero expression and heterochromatin regions. Dashed lines connect observations from the same sample. **(f)** H1.2 ChIP-seq signal at highly expressed enhancers (active enhancers), enhancers with no expression (inactive enhancers), and heterochromatin. For heterochromatin, “TSS” is designated as the center of the heterochromatin interval (Methods). Signals were smoothed for visualization purposes. **(g)** Nucleosome repeat length with respect to the distance from the TSS. NRLs were calculated for each 1,000 bp window with 500bp overlap. The same enhancers and heterochromatin sites were used for (d-g). **(h)** MNase-seq fragment size distributions in H1.2 knockout (blue) and wild-type HeLa cells (orange). **(i)** Nucleosome repeat length phasogram from MNase-seq data. Inset shows linear regression fits for H1.2 knockout and wild-type cells.

We focused on the shoulder region of promoters TSS (±3-5 kb), due to the strong, consistent patterns of correlation observed in this region that are distinct from patterns at the TSS. We fit our model to the observed correlation patterns at these shoulder regions (Methods). The simulation with the best-fit parameter set closely matched the observed periodic patterns of expression correlation across fragment sizes, with a Pearson correlation of 0.9 (**Fig. 3b**). The inferred NRLs were 166 bp for active chromatin and 184 bp for inactive chromatin (**Fig. 3c**). These predictions are in line with our own and other’s observations showing greater nucleosome spacing in active vs. inactive chromatin. For example, analysis of NRL near active enhancer sites in deep-WGS cfDNA from cancer-free volunteers (N=5)^17,47^ showed median NRL of 170 bp (**Fig. 3d, e**, and **Fig. S4**), similar to the ∼179 bp NRL observed using MNase-seq in human CD4+ T cells at H3K27ac-marked regions^34^. Inactive enhancers and heterochromatin in these same samples showed NRLs of 186.5 bp and 187 bp, respectively. These values closely match heterochromatin with the H1 linker protein (184-192bp)^48^. Interestingly, the presence of H1 linker protein is known to increase the effective size of nucleosomes by protecting a small additional amount of DNA in addition to increasing the NRL^49^. The cfDNA fragment sizes predicted by our simulation approach for active and inactive chromatin were 150 bp and 164 bp, respectively, which plausibly correspond to cfDNA with and without H1 linker protein. These observations support the biological validity of the model underlying our simulations and led us to further consider the contribution of the histone H1 protein to the fragmentation patterns we observed.

Given that the H1 linker histone is depleted at active regulatory elements^45,46,50^, we considered whether the reduced effective nucleosome size in these regions might be the result of H1 loss. We analyzed H1 occupancy around active and inactive enhancer groups and found a striking similarity in the profiles of NRL and H1 occupancy near enhancer TSSs. Specifically, as we approached the enhancer TSS, we observed a progressive decrease in H1 occupancy, accompanied by a corresponding shortening of the NRL (**Fig. 3f, g, and Fig. S5**). These results corroborate findings that shorter NRLs are associated with decreased H1 binding^44^. These observations suggest that differences in chromatin fiber structure, defined by nucleosome spacing and linker histone occupancy, may contribute to cfDNA fragmentation signatures associated with transcriptional activity.

To test if H1 linker histone influences nucleosome positioning and fragmentation patterns, we performed MNase-seq on HeLa cells with and without knockout of H1.2. The absence of H1.2 resulted in an overall shortening of fragment lengths, particularly at dinucleosome-sized fragments (**Fig. 3h, S7).** H1.2 knockout cells showed shorter nucleosome repeat lengths compared to wild-type cells, particularly in heterochromatin regions (NRL = 191 bp vs. 195 bp, respectively; **Fig. 3i)**. NRL differences were less pronounced in euchromatin regions, where H1 is already depleted in wild-type cells (NRL = 189 bp vs. 192 bp, **Fig. S7**). This differential effect suggests that H1.2 depletion preferentially shortens NRL in heterochromatin regions where H1 is typically abundant. These results support a model in which removal of H1 near active regulatory elements causes tighter nucleosome spacing, resulting in distinctive DNA fragmentation patterns. These observations suggest that differences in composition of the chromatin fiber, defined by nucleosome spacing and linker histone occupancy, shape cfDNA fragmentation signatures associated with transcriptional activity.

### cfDNA fragmentation near enhancers identifies cancer gene expression from low coverage sequencing data

Having examined a mechanistic basis for the TAS, we tested whether this score identifies activity of enhancers that are context and cell-type-dependent. First, we asked whether the TAS at the enhancers expressed in the LNCaP prostate cancer cell line^51^ correlates with tumor fraction in LP-WGS of cfDNA (median coverage: 0.55x [0.29–0.96]) from patients with metastatic prostate cancer^52^. Despite the small number of informative cfDNA fragments (median 67,531 fragments for each sample across 2,173 enhancers), we observed a strong positive correlation between TAS at prostate-specific enhancer TSSs (normalized by TAS at active blood enhancers; Methods) and cfDNA tumor fraction (r = 0.85, p = 9.2 x 10^-25^; **Fig. 4a**). This result indicates that the TAS can measure expression of enhancers that are specifically active in cancer and not the non-malignant cells that contribute to cfDNA.

**Figure 4.**
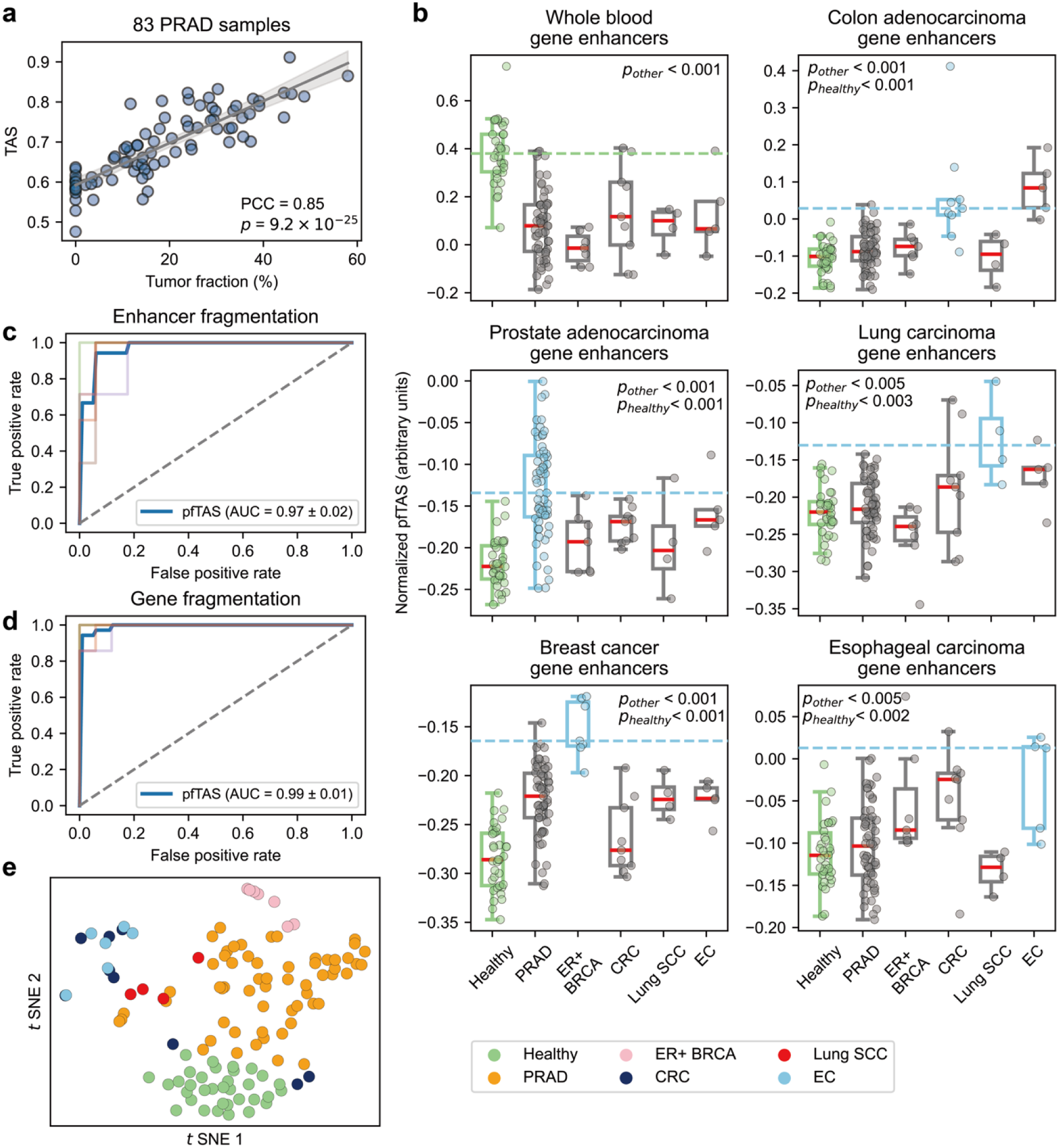
Predicted enhancer activity from low coverage WGS enables cancer detection and typing. **(a)** Ratio of transcriptional activation score (TAS) at prostate cancer-associated eRNAs^51^ to blood-associated eRNAs^33^, plotted against tumor fraction. Data are from cfDNA low-pass WGS (median coverage 0.55x)^52^. **(b)** pfTAS at enhancers linked to gene sets with expression in whole blood (top left) or in the indicated cancer types. Boxplots show pfTAS distributions for published LP-WGS data from patients with the indicated cancer type^31^. Boxplots colored in sky-blue indicate sample–gene set combinations where high expression is expected based on tissue of origin; light-green boxplots represent healthy cfDNA as a baseline. Two-sided Wilcoxon test p-values are indicated for comparisons of pfTAS between cancers in which high expression is expected versus pfTAS for other samples. *p_other_* indicates the *p*-value compared to other cancer samples, and *p_healthy_* indicates the *p*-value compared to healthy samples. **(c)** Classification of cancer vs. healthy from LP-WGS data^11, 21^ using a support vector machine with TAS at enhancers associated with genes^53^ in **(b).** Standard deviation of the AUC was calculated by 5-fold cross-validation. Receiver operating characteristic curves for each fold from cross-validation are shown as transparent lines. Blue lines represent the average across folds. **(d)** As in (c), but using the genes rather than the enhancers. **(e)** Concatenated pfTAS (gene fragmentation and enhancer fragmentation level) from 3,859 genes and 45,211 associated enhancers were projected by *t*-SNE. Cancer types were abbreviated as follows: ER+ BRCA : ER-positive breast cancer (n=7), Lung SCC : squamous cell lung cancer (n=4), PRAD : prostate adenocarcinoma (n=59), CRC : colorectal cancer (n=9), EC : esophageal cancer (n=5).

We next sought to investigate whether fragmentation patterns at enhancers reflected activation of lineage-associated enhancers in cfDNA from patients with cancer. Because the TAS incorporates positional information relative to known TSSs – which may be unavailable for enhancers or ambiguous for genes due to alternative promoters – we devised a position-free TAS (pfTAS). The pfTAS measures median correlation with expression for fragments of a given size without considering positional information (Methods). For each gene, we computed the pfTAS across its linked enhancers using enhancer-gene associations from the GeneHancer dataset^53^. We then aggregated these gene-level scores using tissue- and cancer-specific gene signatures derived from GTEx^54^ and TCGA^55^ (Methods), producing one score per gene signature for each sample. In healthy individuals, we observed significantly higher pfTAS for blood-specific gene signatures, consistent with the hematopoietic origin of cfDNA. In contrast, cancer samples showed elevated pfTAS at lineage-matched gene signatures (**Fig. 4b**). Predicting these signatures from enhancer pfTAS captured activity of these signatures to a similar degree as measuring pfTAS at the genes themselves (**Fig. S8**). We used these signatures of predicted gene transcription as input features to train a support vector machine (SVM) that distinguished cancer from healthy samples (AUC = 0.97; **Fig. 4c**). Classification performance was similar when measuring pfTAS at signature genes directly (AUC = 0.99; **Fig. 4d**). Finally, we combined pfTAS at genes and their linked enhancers and projected these features in low-dimensional space, revealing distinct clusters for different cancer types (**Fig. 4e**). These findings demonstrate that enhancer fragmentation patterns can detect lineage-specific regulatory, even in low-coverage data (median 525,701 fragments per sample across 45,211 enhancers).

### cfDNA fragmentation analysis identifies clinically relevant enhancer and gene transcription in cancer

Next, we asked whether TAS and pfTAS could be used to infer clinically relevant enhancer and gene activity. First, we asked whether the pfTAS could distinguish clinically actionable cancer subtypes in deeply sequenced cfDNA enriched by hybrid capture. We focused on non-small cell lung cancer (NSCLC) and small cell lung cancer (SCLC), which cannot be distinguished reliably by imaging and require different treatments. We analyzed a published cohort of hybrid capture cfDNA sequencing data^56^. Limiting to sites overlapping regions with differential H3K27ac signal in each subtype^36^, we calculated the pfTAS at 123 sites with greater H3K27ac in NSCLC and 361 sites with greater H3K27ac in SCLC. The median pfTAS values at NSCLC-specific sites were significantly higher in NSCLC samples compared to SCLC (Wilcoxon *p*-value =1.8 x 10^-9^; **Fig. 5a**). Conversely, SCLC samples demonstrated higher scores than NSCLC samples at SCLC-specific enhancers (Wilcoxon *p*-value =1.5 x 10^-9;^ **Fig. 5b**). The ratio pfTAS at NSCLC vs SCLC-associated enhancers distinguished the two subtypes (AUC = 0.94; **Fig. 5c**). Classification was not solely driven by differences in tumor fraction, because classification based on tumor fraction yielded a lower AUC of 0.88 (**Fig. 5c, S9**).

**Figure 5.**
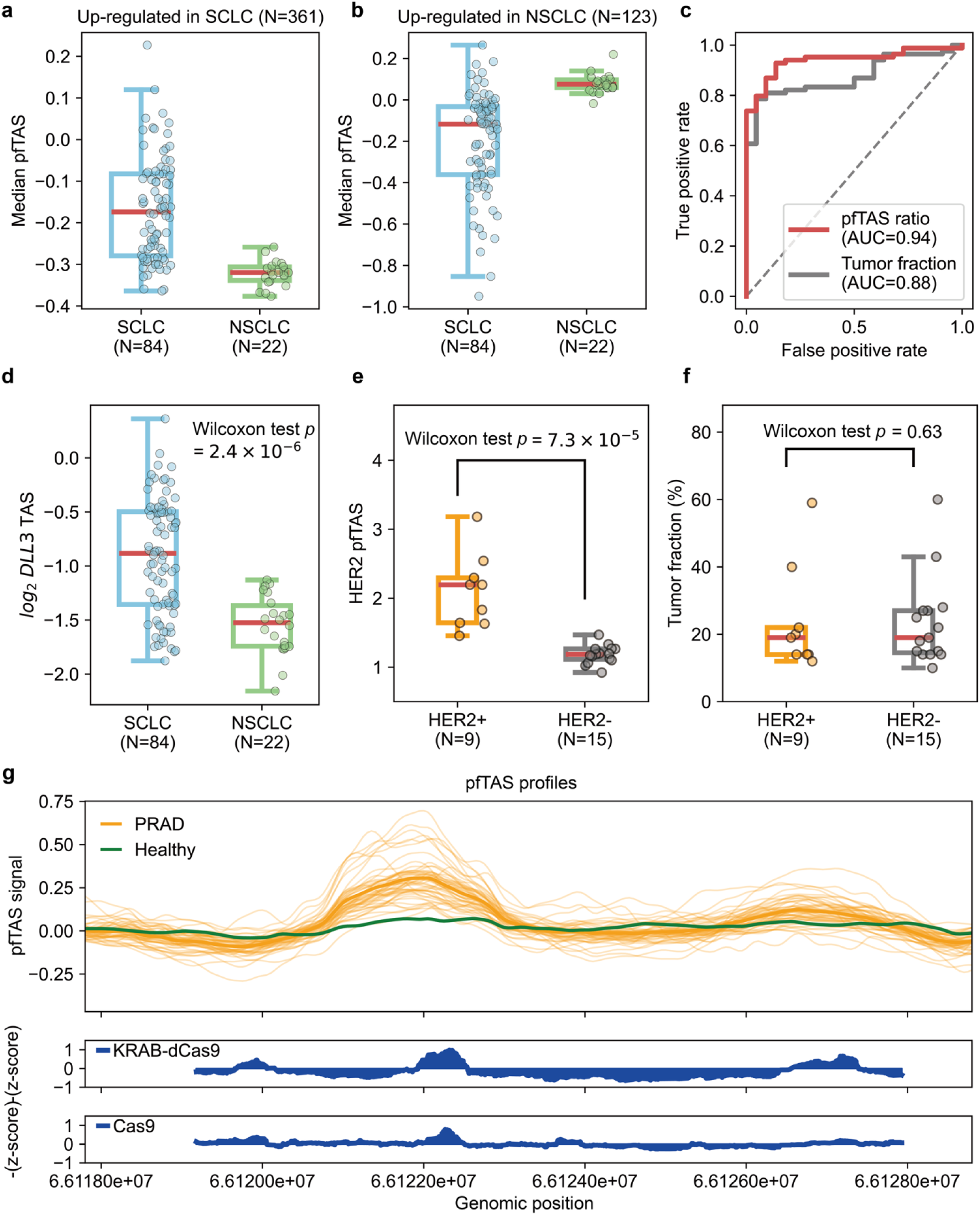
Enhancer and gene fragmentation patterns are preserved in high coverage sequencing data. **(a)** and **(b)** The box plots represent the pfTAS distributions of SCLC and NSCLC, calculated from cfDNA hybrid capture data at N=361 enhancers that are active in SCLC but not NSCLC (median coverage 67.5x) and N=123 enhancers that are active in NSCLC but not SCLC (median coverage 38.9x). **(c)** AUROC curves from pfTAS ratio between SCLC and NSCLC sites and cfDNA tumor fraction to classify SCLC and NSCLC. **(d)** The *DLL3* TAS distributions (median coverage 85.1x) **(e)** The *ERBB2* (HER2) pfTAS distribution. HER2-positive and negative samples were classified by using amplification and Immunohistochemistry (Methods). **(f)** cfDNA tumor fraction distribution of samples in **(e). (g)** TAS scores calculated across the AR enhancer locus for N=49 PRAD samples and 1 healthy sample. Results from a tiling CRISPR/CRISPRi screen^58^ are shown below. Higher (z-score) indicates reduced proliferation of LNCaP cells with suppression of the indicated site.

Subsequently, we asked whether TAS can detect elevated expression of the *DLL3* gene, which encodes a targetable cell surface protein that is expressed in ∼95% of SCLC^57^. We measured the TAS at *DLL3*, centered on the annotated TSS with high H3K4me3 signal from SCLC cfChIP-seq data^36^ (**Fig. S10**). TAS values at *DLL3* were significantly higher in SCLC samples compared to NSCLC (Wilcoxon *p*-value = 2.4 x 10^-6^; **Fig. 5d**). These results show that the cfDNA fragmentation patterns we describe are preserved after hybrid capture enrichment and may distinguish clinically actionable cancer subtypes and reflect expression of therapeutic targets.

To test the generalizability of this approach, we asked whether cfDNA fragmentation patterns could identify expression of another targetable cell-surface protein, HER2. We analyzed cfDNA fragments sequenced with a commercial gene sequencing platform (FoundationOne), comparing 9 patients with HER2+ cancers to 15 patients with HER2-cancers, assessed by the presence or absence of *ERBB2* amplification and concordant HER2 protein expression on immunohistochemistry. We applied the pfTAS due to the presence of multiple annotated TSSs for the HER2-encoding gene *ERBB2* and to maximize coverage across multiple exons. The pfTAS at *ERBB2* were significantly higher in the HER2-positive samples than in the HER2-negative samples (**Fig. 5e**), despite similar tumor fractions between these groups (**Fig. 5f**). This result provides another example of how this approach can infer expression of a targetable gene in cancer.

We next asked whether the pfTAS could detect transcription of a functionally validated enhancer of the androgen receptor gene (*AR*) in deeply sequenced cfDNA from metastatic prostate cancer (median coverage = 116x)^47^. The *AR* enhancer is inactive in adult tissues, but reactivated in prostate cancer as a mechanism of resistance to androgen deprivation therapy^58^. We applied a sliding window across the *AR* enhancer and computed the pfTAS in 49 metastatic prostate cancer cfDNA samples. Regions within the enhancer that exhibited high pfTAS values overlapped with functional elements identified by tiling Cas9 and KRAB-dCas9 screens of the *AR* enhancer region^58^ (**Fig. 5g**). These screens measured the impact of enhancer disruption (Cas9) and repression (KRAB-dCas9) on proliferation of LNCaP, which requires *AR* activity. Prostate cancer cfDNA, but not healthy cfDNA, showed focal elevations of the pfTAS at the functionally important region of the *AR* enhancer locus (indicated by the -z-score). This finding demonstrates that cfDNA fragmentation patterns can identify activity of resistance-associated enhancers in cancer.

## Discussion

This study identified patterns of cfDNA fragmentation that correlate with transcription at enhancers, a close correlate of enhancer activity. We propose a mechanistic basis for these patterns, supported by generative modeling of cfDNA that recapitulates our observations. This model suggests that a simple parameter set that describes the chromatin fiber structure – nucleosome repeat length and effective nucleosome size – can account for the systematic differences in cfDNA fragmentation at active vs. inactive TSSs. In deeply sequenced cfDNA, we validated key predictions of our model, including tight nucleosome spacing near active TSSs and looser spacing at inactive TSSs. We defined a transcriptional activation score (TAS) based on these cfDNA fragmentation patterns and demonstrated its ability to non-invasively detect tumor-specific gene expression signatures and differentiate between cancer types in low-coverage cfDNA data. Importantly, we show these patterns are preserved with hybrid capture, enabling targeted deep sequencing to predict the expression of biologically important resistance enhancers (e.g., the *AR* enhancer) or therapeutic targets (e.g., DLL3 and HER2) from cfDNA.

Our model provides a unifying framework for cfDNA fragmentomic features associated with chromatin structure and reconciles apparently contradictory observations from prior studies. It explains the observation that both very short and long fragments are associated with active chromatin^19,20^, as well as complex biases in the size distributions of H3K27ac-marked regulatory elements^12^. Our model also explains a recent finding that nucleosome repeat length inferred form cfDNA is wider at genes with low expression^59^. Strikingly, a substantial proportion of fragments have sizes that are significantly correlated with enhancer expression after FDR correction (14.8% of fragments within an 8kb of the TSS of enhancers, 25.9% for promoters), but the magnitude and direction of correlation depend on distance from the TSS. While we detected increased fragmentation entropy precisely at the TSS, our model suggests that differences at the TSS “shoulders” in cfDNA fragment sizes can be accounted for by consistent (non-random) changes in nucleosome position and effective size. Entropy signals may reflect a combination of nucleosome depletion around the TSS (as previously proposed) as well as distinctive fragment lengths that arise due to closely spaced nucleosomes near active TSSs. Finally, our findings may account for the locus-specific increased variance in fragment length observed in cancer^3^. This variance could arise from fragmentation of active and inactive chromatin at genes with differential expression in cancer vs. non-malignant cells contributing to cfDNA.

The model also captures the surprising, yet biologically consistent, pattern of nucleosome spacing and cfDNA cleavage rates increasing in regions of heterochromatin and decreasing in euchromatin near active TSSs. This finding may seem counterintuitive given the association of accessible chromatin with active regulatory elements. While the TSS itself at active enhancers shows accessibility and nucleosome depletion, immediately adjacent chromatin is comprised of tightly spaced nucleosomes. This tighter spacing results in a lower propensity to cleave between nucleosomes and a higher portion of dinucleosome fragments in cfDNA. These findings are consistent with early studies of nucleosome positioning using MNase-seq, which noted tighter spacing in euchromatin compared to heterochromatin^34^.

Our model is supported by our analysis of HeLa cells with H1.2 knock-out, as well as recent studies of chromatin fiber structure and its relationship to gene expression, NRL, and the presence of the H1 histone protein. The density of H1 has been proposed to regulate local chromatin structure^46,60,61^. Structural studies show that H1 presence is closely associated with NRL and is important for determining the structure of the chromatin fiber and chromatin compaction^44,62^. Previous studies have shown that H1 is enriched in heterochromatin and depleted in euchromatin, particularly around gene TSSs^46,63^, in agreement with our results. This relationship may be causal, wherein H1 depletion decreases NRL *in vitro*^50^ and *in vivo*^64^ and increases expression of noncoding RNAs^50^. The size and NRL parameters we identified for inactive vs. active chromatin are close to reported values for chromatin fibers with vs. without H1 linker protein. H1 suppresses nucleosome unwrapping and decreases exposure of DNA to nuclease activity, accounting for the longer effective nucleosome size in inactive chromatin where H1 localizes^65^. Our results suggest that complex associations between cfDNA fragment lengths and transcription may be explained by differential representation of H1. We propose a model where H1 is depleted in active chromatin near expressed TSSs, resulting in smaller effective nucleosome sizes, tighter spacing, higher D/M ratio in cfDNA, and the periodic correlation we observe between fragment size and expression.

This study has several limitations. The effect of H1 depletion on NRL we identified was modest, potentially due to other determinants of NRL and because of redundancy between H1 linker variants. Additional mechanistic studies are warranted to define the influence of each linker histone variant on cfDNA fragmentation. In addition, future studies will be needed to determine if cancer-specific fragmentation patterns are determined by differences in H1 occupancy, an intriguing possibility given the observed suppression of H1 in cancer^66^. An additional limitation is that enhancer fragmentation signatures from LP-WGS data did not fully resolve distinct cancer types. This is likely due to the low depth of coverage in the data analyzed here (<1x coverage). Future studies should explore how sequencing depth affects the ability to resolve activation of individual enhancers and enhancer modules. It will be necessary to test how predictions of enhancer activity in cancer are influenced by tumor fraction.

Inferring enhancer expression from cfDNA is a novel approach to studying gene regulation from plasma that has several advantages. Enhancers are closely linked to cell types and states and their activity can inform disease biology. Many enhancers are co-regulated, enabling aggregation across enhancer modules to overcome sparsity in low-coverage sequencing data. In addition, this approach has several advantages over existing methods for studying gene regulation in cancer, such as DNA methylation^67^ and hydroxymethylation^68–70^ profiling or cfChIP-seq. Because it requires only sequencing data, the TAS and pfTAS can be applied to existing data from cfDNA sequencing platforms, including hybrid capture data from a widely used commercial sequencing platform. It could be incorporated into platforms assessing mutations with deep sequencing, to provide both genetic and gene-regulatory landscapes of cancer as it evolves in response to treatment. This approach could aid treatment selection by predicting expression of targets for a large and growing class of treatments targeting cell surface proteins in cancer, such as antibody drug conjugates, T-cell engagers, and CAR T-cells. Finally, our observations should enable studies of the *epigenetic* drivers of response and resistance in human cancers at scale, which have been poorly characterized compared to genetic mutations.

## Acknowledgments

This work is supported by the Fund for Innovation in Cancer Informatics, the Wong Family, the Damon Runyon Cancer Research Foundation, the US Department of Defense (DoD) (award W81XWH-21-1-0358), the Prostate Cancer Research Foundation, and the National Institute of Health (NIH) (1U01CA296432-01). D.F. is supported by grant T32HG002295 from the NIH. G.S.G. is supported by an NIH T32CA009172 award from the National Cancer Institute (NCI). J.E.B. is supported by the US DoD (W81XWH-20-1-0118, HT9425-23-1-0048).

## Methods

### DNA sequence data processing

Paired-end DNA sequencing data were aligned to the hg19 reference genome after adapter trimming with fastp (0.23.2)^71^. Burrows–Wheeler Aligner (BWA) (0.7.17)^72^ was used for alignment, with insert size modeling disabled. Reads with any of the following were removed by Samtools (1.11)^73^: (1) mapping quality (MAPQ) less than 5, (2) no proper read pair, or (3) a read pair mapped to a different chromosome. Finally, the filtered reads were converted to fragment locations in BEDPE format using bedtools (2.30.0)^74^. Fragments that shared the same start and end position were considered likely PCR duplicates and collapsed into a single fragment. All analyses of coverage used fragment locations in BEDPE format, so that any unsequenced DNA between read pairs contributed to coverage. High noise regions^75^ were excluded from all subsequent analysis.

For the analysis of the *ERBB2* locus, BAM files from FoundationOne liquid biopsy test aligned to the hg19 human genome build were provided by Foundation Medicine. These tests were ordered as part of the clinical care of patients at Dana-Farber Cancer Institute, and the sequencing results were returned to DFCI and analyzed under an IRB-approved protocol. The deeply sequenced WGS data from Herberts *et al.*^47^ (two healthy samples used for the NRL analysis in **Fig. 3e** and **Fig. S4**, and 49 PRAD samples and one cancer-free control used for the AR enhancer visualization in **Fig. 5e**) were obtained as bam files aligned to the hg38 reference genome and filtered further. We noted that reads from certain sequencers used to generate these data produced shorter fragment size distributions compared to the remaining sequencers, even when comparing fragments from the same sample. Therefore, we discarded reads with sequencer IDs HISEQ101_212, HISEQ101_213, HISEQ102_181, HISEQ102_197, HISEQ103_173, HISEQ103_196, HISEQ104_171, HISEQ104_177, HISEQ104_178, HISEQ104_224, HISEQ105_221. To avoid the need for realignment, we lifted over genomic annotations (*e.g.* TSSs) to the hg38 reference genome for analyses involving these samples.

### Analysis of cfDNA fragment sizes segregated by chromatin states

To define consensus chromatin states, we downloaded the 27 ChromHMM segmentation files for hemtatologic cells from the Roadmap Epigenomics Project^24^. These files divide the genome into 200 bp bins and assign a chromatin state label to each bin using for each sample using ChromHMM^35^. To identify consensus states, we retained only those bins for which at least 50% of the samples were labeled with the same chromatin state. All other bins were excluded from the analysis. The D/M Ratio was defined as the ratio of dinucleosome-sized fragment (300-340 bp) counts to mononucleosome-sized fragment (140-180 bp) counts. Dinucleosome modal size is defined as the most common size of fragments between 300-340 bp.

### Correlation analysis of expression with cfDNA fragment size and location

cfDNA fragmentation patterns – represented by the sizes of fragments and their distance upstream or downstream from transcriptional start sites (TSSs) – were analyzed for correlation with eRNA expression. eRNAs were divided into 30 quantiles based on their expression levels measured by PRO-cap sequencing in lymphoblastoid cell lines^33^. For each quantile, cfDNA fragments were grouped into 5-bp size bins (50–55 bp, 55–60 bp, …, 496–500 bp) and 50-bp distance intervals spanning −8 kb to +8 kb around the TSS. Within each quantile, we calculated the normalized density of fragments in each bin, defined as the number of fragments in the bin divided by the total number of fragments in the quantile. For each bin, the vector of 30 fragment densities (one per expression quantile) was correlated (Pearson’s r) with the corresponding median expression values across quantiles. Correlation heatmaps were generated separately for enhancers and promoters, showing correlations of cfDNA fragment densities across fragment size and distance-to-TSS bins. For promoters, TSSs were stratified by RNA-seq expression levels in peripheral blood mononuclear cells (PBMCs) from healthy individuals^14^ using the same quantile approach.

### Transcription factor enrichment analysis

Fragments from 34 pooled healthy cfDNA samples^31^ within ±500 bp of enhancer TSSs (defined in lymphoblastoid cell lines) were analyzed for the top 10,000 enhancers ranked by PRO-cap expression levels. The genomic sequences corresponding to the long and short fragments were extracted and analyzed using HOMER^76^ (findMotifsGenome.pl, with the parameters *-size given -mask -p*) to identify enriched transcription factor motifs. The significance of motif enrichment reported by HOMER was compared for long vs. short fragments.

### Transcriptional Activation Score (TAS) calculation

The Transcriptional Activation Score (TAS) is calculated by summing cfDNA fragment counts at tiled genomic positions relative to a TSS where expression is correlated with fragment size and location (see **Fig. 2a**). Genomic positions within 5kb upstream or downstream of a TSS were grouped in 5bp tiles, and fragment lengths were binned by 5bp. The TAS is derived by dividing the summed fragment counts in positively correlated bins by those in negatively correlated bins.

The formula is expressed as:

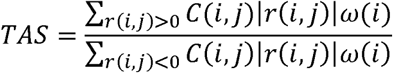

where, *i* and *j* indicate distance from the TSS and fragment size, respectively. *C(i,j)* represents the number of a *j*-size fragments in the *i* position. *r(i,j)* represents smoothed PCC of the corresponding *i,j* position (**Fig. 2a**; Supplementary tables 1 and 2). PCC was smoothed by a Gaussian filter with sigma equal to 7.5 bp for fragment size and 75 bp for distance. Bins with values of *r(i,j)* < 0.2 were omitted to focus on bins with high correlation (**Fig. S2**) w(i) indicates weight function for the position. The w(i) is defined by a Gaussian function with a mean of 0bp and sigma of 1,000 bp to give regions closer to the TSS higher importance. For the pfTAS calculation, *C(i,j)* was substituted with the median value across all distances from the TSS within 5kb, and the weight function for the position was set to 1. *C(i,j)* values for enhancers and promoters are listed in Supplementary Tables 1 and 2. Source code for TAS is available in our laboratory GitHub repository (https://github.com/Baca-Lab/TAS_fragmentomics)

### Modeling nucleosome size and NRL

We built generative models of cfDNA fragmentation to identify parameters that best fit the observed correlations of expression with fragment size and distance from TSS. The corresponding notebook is uploaded to our laboratory GitHub repository at https://github.com/Baca-Lab/TAS_fragmentomics/blob/main/simulations.ipynb.

We posited that differences in cfDNA fragment length distributions at “active” vs. “inactive” chromatin can be explained by variation in two simple parameters: nucleosome repeat lengths (NRLs) and nucleosome footprint sizes (corresponding to the amount of DNA protected from nuclease digestion due to contact with histone proteins). We assumed that these two features have distinct but partly overlapping distributions in active vs. inactive chromatin. We define parameter set {*NRL_active_*, *NRL_inactive_*, *size_active_*, *size_inactive_*} to represent the means of normal distributions for these features in active vs. inactive chromatin.

To fit these parameters to observed data, we simulated nuclease digestion of DNA bound to nucleosomes. Nucleosome positions were simulated along 100kb of DNA. Each simulation tested mean NRL and size values for active chromatin { *NRL_active_*, *size_active_*,} and for inactive chromatin {*NRL_inactive_*, *size_inactive_*}, then assigned nucleosome positions and sizes by drawing from normal distributions with these means. Standard deviations for NRLs, nucleosome sizes, and spacing between nucleosomes were set at 10bp. For each simulation, approximately 10,000 cuts were attempted at random positions. If a cut attempt landed at a coordinate not protected by a nucleosome, the DNA was cut accordingly. If a cut attempt occurred where a nucleosome was present, a cut was generated with 0.001 probability.

*NRL_active_test_ =* seq(140, 215, by = 2)

*NRL_inactive_test_ =* seq(140, 215, by = 2)

*size_active_test_ =* seq(120, 190, by = 2)

*size_inactive_test_ =* seq(120, 190, by = 2)

Combinations of parameters where active chromatin had larger NRLs or protection size than inactive chromatin were excluded from the search for efficiency. This constraint reflects prior observations that active chromatin exhibits a shorter NRL compared to inactive chromatin and that the depletion of histone H1 in active chromatin leads to reduced nucleosome protection compared to inactive chromatin.

To generate the correlation profile observed between fragment size and gene expression, we averaged the correlations between cfDNA fragment density and gene expression across distance bins between 3 to 5 kb from the TSS, excluding the central ±3 kb window.

For each parameter set, we performed multiple independent simulations of DNA fragment lengths from 50–500 bp and aggregated them into 5 bp bins for active and inactive chromatin separately. For each bin, we calculated the ratio of fragment density in active chromatin to inactive chromatin, adding a small term to avoid division by zero. These ratios were log₂-transformed to generate a simulated log-ratio profile describing the fragment size-dependent bias toward active vs. inactive chromatin. We computed the Pearson correlation between this simulated profile and the observed fragment size–expression correlation profile.

We used Bayesian optimization with Expected Improvement acquisition to identify optimal parameters. The objective minimized 1 + r, where r is the mean Pearson correlation across batches. Five independent runs (200 evaluations each, starting from 20 uniformly sampled points) converged to consistent estimates for optimal parameters (*NRL_active_max_, NRL_inactive_max_*, *size_active_max_, size_inactive_max_*) that maximized agreement between simulated and observed patterns. These parameters describe the NRL and size distributions that are *most differential* between active and inactive chromatin.

We then returned to the calculation of *NRL_active_*, *NRL_inactive_*, *size_active_*, and *size_inactive_*. We assumed that effective nucleosome size and NRL are drawn from normal distributions that partly overlap between active and inactive chromatin. We interpreted the Bayesian-optimized values of {*NRL_active_max_, NRL_inactive_max_*} and {*size_active_max_, size_inactive_max_*} as the NRLs and fragment sizes that are the most different between the distributions represented by means {*NRL_active_*, *NRL_inactive_*} and {*size_active_*, *size_inactive_*}. Therefore, to identify {*size_active_*, and *size_inactive_*}, we fit partially overlapping normal distributions to the top Bayesian-optimization solutions for active and inactive nucleosome footprint size. To ensure that distributions are partly overlapping, we required the means *size_active_*, and *size_inactive_*to be within three standard deviations of each other. Finally, we selected {*size_active_*, *size_inactive_*} where the maximum ratio of active to inactive fragment densities is closest to *size_active_max_* and the minimum of this ratio is closest to *size_inactive_max_*. We followed an identical procedure to identify optimal values of *NRL_active_* and *NRL_inactive_*.

### Nucleosome spacing analysis and H1 ChIP-seq analysis

To estimate nucleosome spacing patterns around genomic regions of interest, we used deeply sequenced WGS data from five cancer-free volunteers (two from Snyder et. al^17^; the other three from Herberts et. al^47^) to calculate high-resolution nucleosome spacing patterns (35 bp windows, tiled in increments of 5bp around the TSS). We counted the number of fragments spanning the window vs. ending in the window to calculate the nucleosome footprint score (NFS) as described^77^. Nucleosome repeat lengths were calculated using NucTools^78^ with the filtering threshold parameter increased to 1,000 to account for the high depth of sequencing. Active enhancers and inactive enhancers were defined by their mean expression^33^. The top 5,000 enhancers were defined as active enhancers, and the bottom 5,000 enhancers were defined as inactive enhancers. To define heterochromatin sites for comparison, we identified all consecutive intervals of ChromHMM “Heterochromatin” consensus annotations and took the longest 5,000 such intervals. The “TSS” was arbitrarily assigned as the center of each heterochromatin interval. Wig files containing H1 ChIP-seq signal (control subtracted) were obtained from a published study and used without further processing^79^. For display, H1 ChIP-seq signal was averaged in 10bp bins.

### Chromatin and RNA profiling of HeLa wild-type and H1.2 knockout cells

HeLa wild-type (Abcam, ab255928) and H1.2 knockout (Abcam, ab261794) cells were cultured in high-glucose DMEM supplemented with 10% fetal bovine serum and 1% penicillin-streptomycin. Knockout validation was confirmed by Western blot using anti-H1.2 antibody (Abcam, ab4086) and with anti-β-actin antibody (Invitrogen, MA1-140) as a loading control **(Fig. S6)**. For nucleosome profiling, nuclei were isolated and digested using the EZ Nucleosomal DNA Prep Kit (Zymo Research, D5220) with Atlantis dsDNase. Digestions were performed with 0.5 units for wild-type and 0.35 units for H1.2 knockout cells for 30 min at 42 °C, followed by nucleosomal DNA purification according to the manufacturer’s protocol. Sequencing libraries were constructed from the digested DNA using the KAPA HyperPrep Kit (Roche, KK8504). RNA was extracted from parallel cultures using the RNeasy Mini Kit (Qiagen, 74134). Nucleosome DNA and RNA were both sequenced on the Illumina NovaSeq platform (Novogene).

Euchromatin and heterochromatin regions were defined by aggregation of active chromatin states (TssAFlnk, TxWk, Enh, TssA, TxFlnk, EnhG, Tx) and inactive chromatin states (ReprPC, BivFlnk, Quies, ZNF/Rpts, ReprPCWk, Het, EnhBiv) from ChromHMM annotations to overcome data sparsity. NRL profiles for each group were computed using NucTools^78^ with the default filtering threshold (20).

### Lineage-associated gene and enhancer analysis

Lineage-associated genes were defined using processed RNA-seq data from TCGA^55^ and GTEx^54^. TCGA RNA-seq data were retrieved with TCGAbiolinks^80^ in the form of TPM. TPM values for gene expression in whole blood were downloaded from GTEx portal. For each cancer type, protein-coding genes satisfying the following criteria were defined as lineage-associated genes: (1) *p*-value < 0.01 between samples of one cancer type and all other cancer types (Wilcoxon rank sum test) and (2) mean fold change > 2. Finally, genes identified as lineage-associated in more than one cancer type were discarded. Whole blood cell-associated genes were defined analogously, and the top 600 genes by TPM fold-difference between whole blood and all other tissues were selected. Enhancer regions were linked to the lineage-associated genes using the GeneHancer database^53^, retaining only enhancer–gene associations high confidence (score > 10).

pfTAS were calculated for each gene level. For the gene fragmentation level, genomic region from upstream 8kb to gene body was used to calculate pfTAS. For the enhancer fragmentation level, if multiple enhancers were linked to one gene, such enhancers were aggregated to calculate a single pfTAS value for the gene. pfTAS values were standardized for each sample. To filter out noisy pfTAS values due to low coverage, we only included genes with ≥ 20 fragments across associated enhancers in all samples. For the gene fragmentation analysis, similarly, genes with ≥ 20 fragments from 8kb upstream through the gene body were included. GENCODE v19^81^ annotations were used to define genomic region for each gene.

cfDNA samples were represented as vectors of pfTAS across lineage-specific genes. For binary classification of cancer vs. healthy samples, support vector machines (SVMs) with radial basis function (RBF) kernels were trained on these vectors, with performance evaluated via 5-fold stratified cross-validation. Receiver operating characteristic (ROC) curves were generated by averaging across folds.

For visualization, *t*-distributed stochastic neighbor embedding (*t*-SNE) was performed on a matrix of samples by pfTAS with Principal Component Analysis (PCA) initialization. pfTAS values for gene bodies and associated enhancers were concatenated prior to *t*-SNE projection for this visualization.

### Hybrid capture (targeted sequencing) data analysis

SCLC- and NSCLC-specific regulatory elements were defined in a previous study^36^ based on differential H3K27ac ChIP-seq signal between these two cancer types. From these differential sites, we selected those with high coverage in the cfDNA hybrid capture dataset from Hiatt et al.^56^. All SCLC and NSCLC cfDNA datasets that passed quality checks from the original paper^56^ were included regardless of their cfDNA tumor fraction estimation. We retained sites where all samples had coverage > 50% of the reported sample-specific mean target coverage. pfTAS were calculated for each site and standardized with a mean of 0 and standard deviation of 1 for each sample. For pfTAS ratios, values were used without standardization because sample-specific variation captured in the numerator and denominator was expected to cancel out. H3K4me3 ChIP-seq coverage profile were retrieved from the El Zarif et al. ^36^. *DLL3* TAS values were normalized by median TAS values of previously defined housekeeping genes^82^. Housekeeping genes were included for normalization if they had sufficient coverage, defined as > the mean sample-specific target coverage in > 90% samples. For FoundationOne data used in the HER2 analysis, the samples with *ERBB2* amplification identified by FoundationOne and HER2 protein expression on immunohistochemistry (IHC) analysis were classified as HER2 positive. Other samples with no reported *ERBB2* amplification and absent HER2 staining on IHC were classified as HER2 negative. Subsequently, the *ERBB2* pfTAS was calculated for each sample and normalized by pfTAS signal at housekeeping genes^82^.

### AR enhancer pfTAS analysis

pfTAS scores were computed in 3 kb sliding windows with a 50bp step, across the *AR* enhancer region (hg19 coordinates chrX:66,116,500–66,133,950). Windows with fewer than 150 overlapping cfDNA fragments were set to zero to minimize noise from low-coverage regions. To enable comparison across samples, pfTAS profiles were median centered by subtracting the median pfTAS across the region for the corresponding sample.

