## Supplementary Figures S1-S11 for "A chromatin fiber model explains cell-free DNA fragmentation signatures of active regulatory elements"

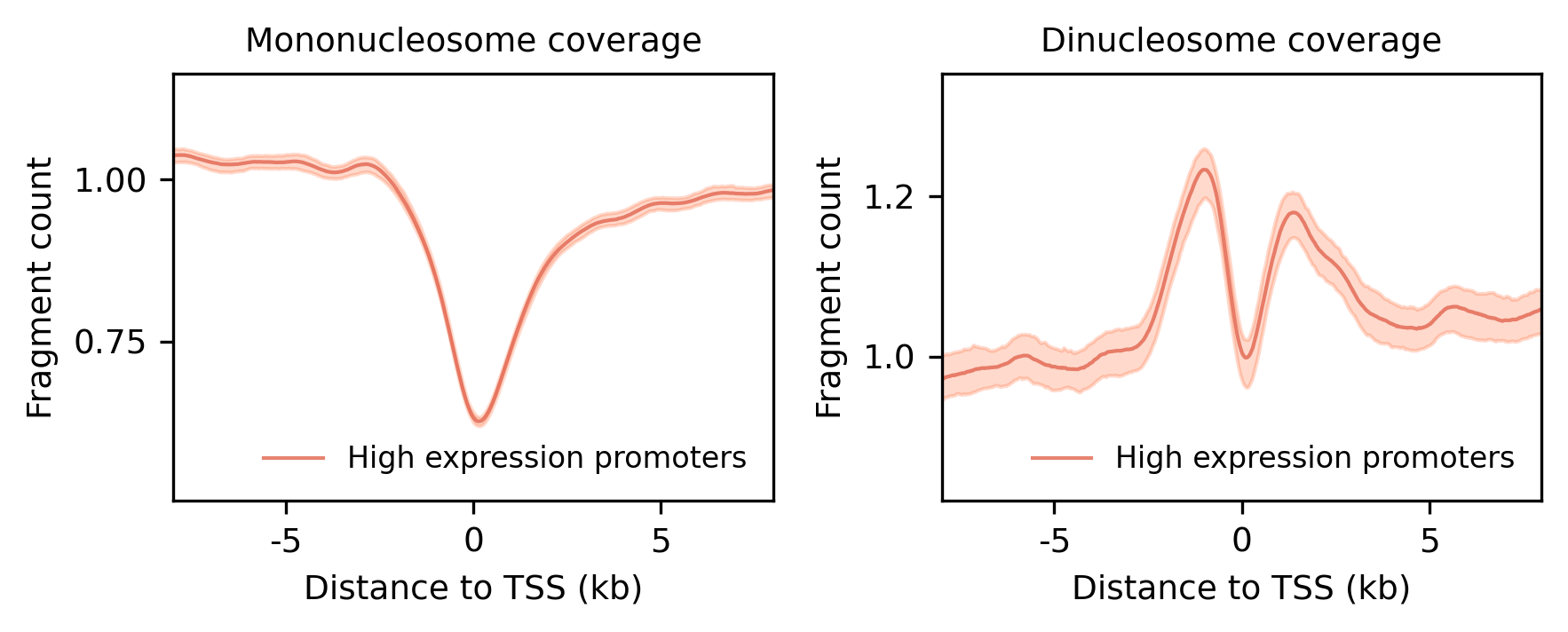

**Figure S1. Fragment midpoint coverage near promoter TSSs as in Fig. 1c, d.** Aggregated midpoint count of mononucleosome size (140-180 bp) and dinucleosome size fragments (300-340 bp) near gene the transcriptional start sites (TSS) of highly expressed genes**.** Shaded regions represent 95% bootstrap confidence intervals.

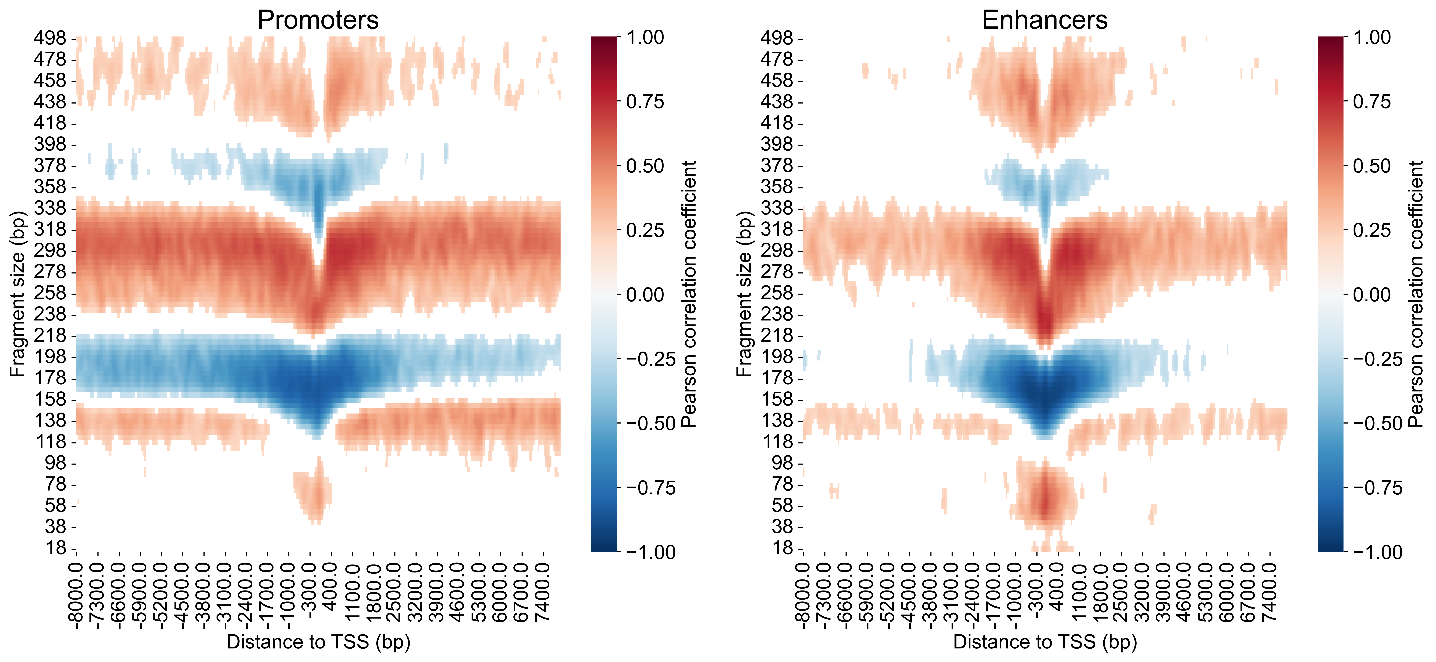

**Figure S2. Heatmap showing correlation values used in the transcriptional activation score (TAS) calculation.** The heatmaps show the Pearson correlation coefficient (PCC) between gene or enhancer expression and the relative abundance of fragments for each distance and fragment size pair. The pairs of fragment sizes and distances to TSS with | PCC | < 0.2 were masked (colored white in the figure), and all other pairs were used for TSS calculation. The correlation matrix was smoothed using Gaussian filter.

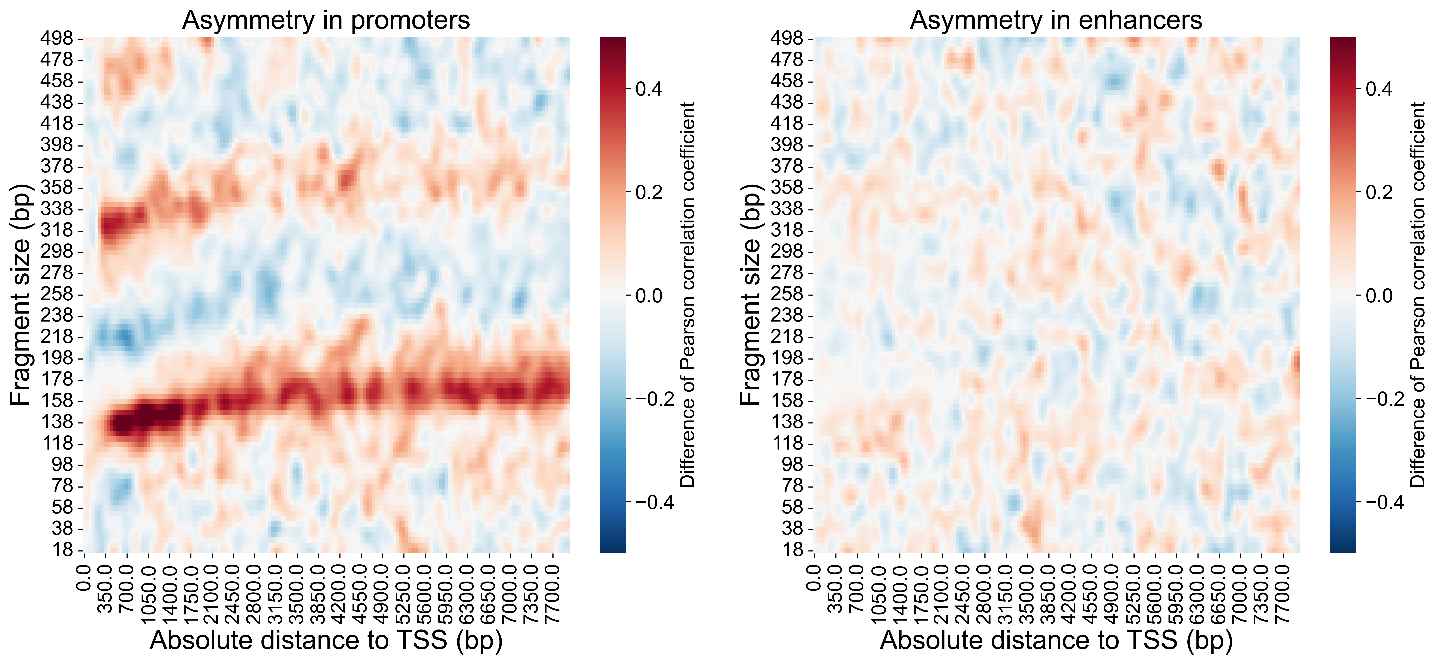

**Figure S3. Symmetry analysis of the expression correlation heatmap.** Heatmaps showing correlation of expression with fragment size and location (see Fig. 2a and e) were assessed for symmetry around the TSS. Correlation values for each heatmap tile downstream of the TSS were subtracted from the corresponding values upstream of the TSS. These correlation differences are displayed and clipped at $\pm$ 0.4 for visualization purposes.

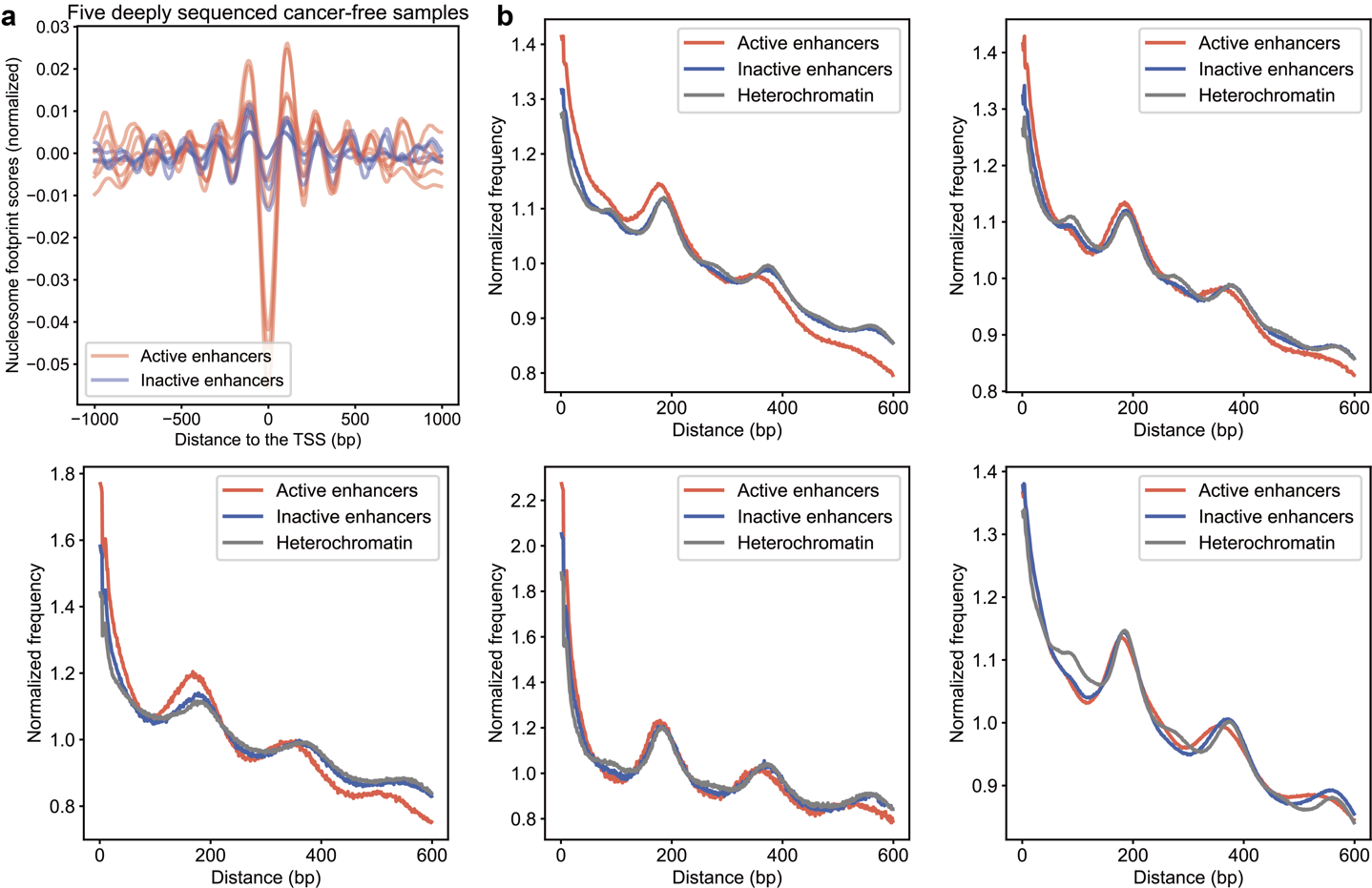

**Figure S4. Nucleosome repeat lengths vary with chromatin state. (a)** Nucleosome footprint score (NFS)^76^ profiles at active and inactive enhancers in five deeply sequenced cancer-free cfDNA samples^17,47^ NFSs were mean-centered for visualization purposes. (**b)** Phasograms^77^ showing the distribution of distances between nearby fragments used for NRL calculations are presented for each of the five cfDNA samples. The phasograms were separately calculated for fragments overlapping active enhancers (top 5,000 by expression in lymphoblastoid cell lines^33^), inactive enhancers (5,000 with no expression), and heterochromatin regions based on consensus chromHMM annotations (see Methods).

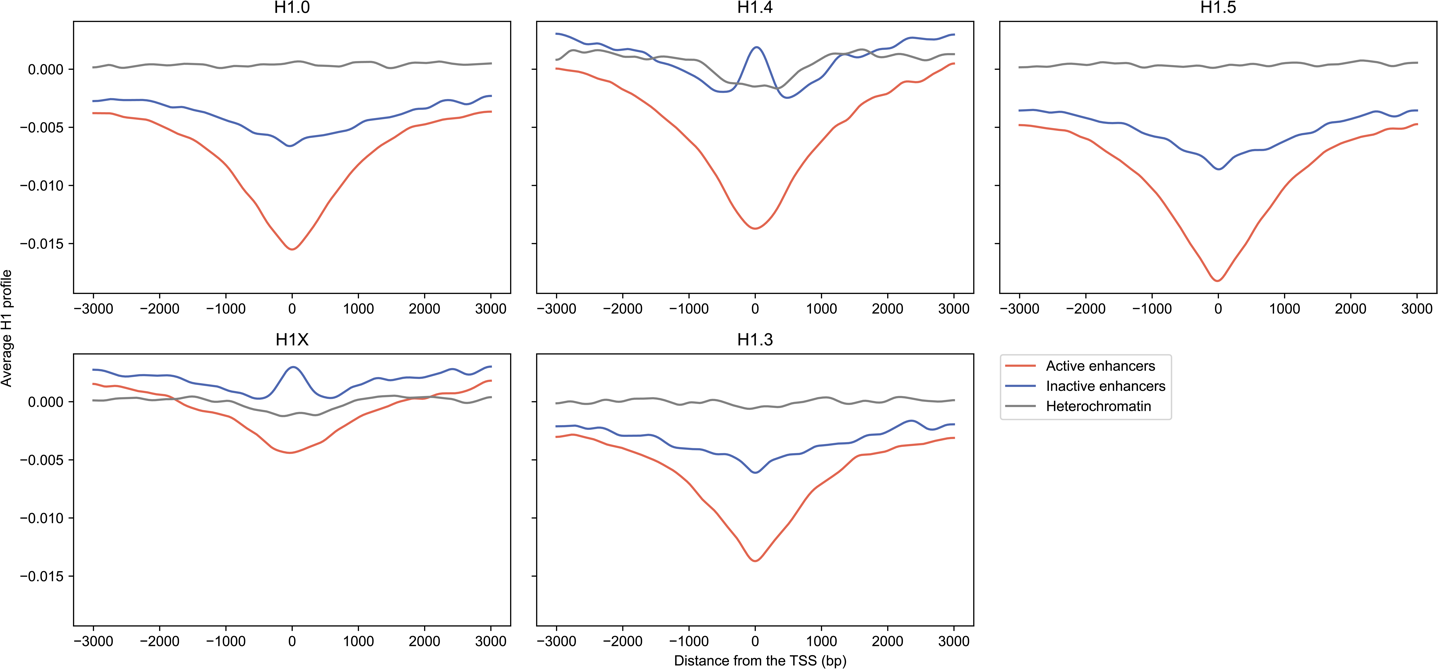

**Figure S5. H1 variant abundance near TSS.** Mean coverage from published histone H1 variant ChIP-seq datasets^78^ versus distance from enhancer TSSs. ChIP-seq coverage is normalized to input controls and obtained from wig files from the published study^78^. Active and inactive enhancers were defined by their eRNA expression in lymphoblastoid cells, as in Fig. 3. Heterochromatin sites are based on consensus chromHMM annotations in blood cells, as in Fig. 3. Signals were smoothed for visualization purposes.

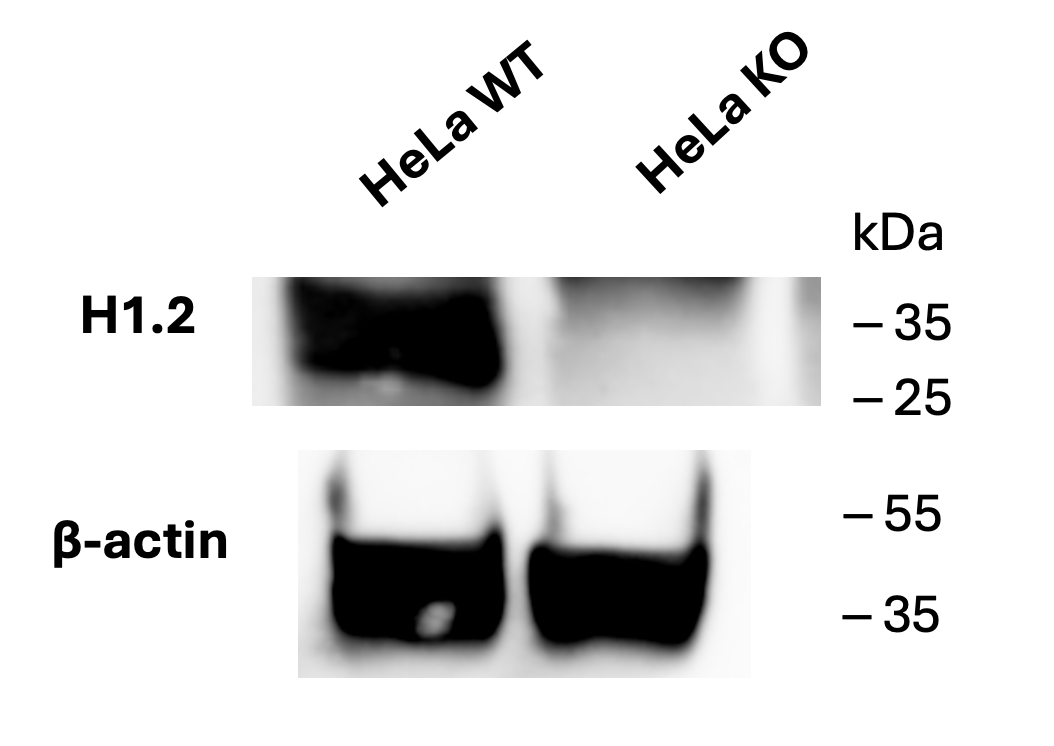

**Figure S6. Validation of histone H1.2 knockout in HeLa cells by Western blot analysis.** Western blot showing Histone H1.2 protein expression in wild-type (WT) and histone H1.2 knockout (KO) HeLa cell lines. The H1.2 band (~35 kDa) is present in WT cells but absent in KO cells. β-actin (~55 kDa) serves as a loading control.

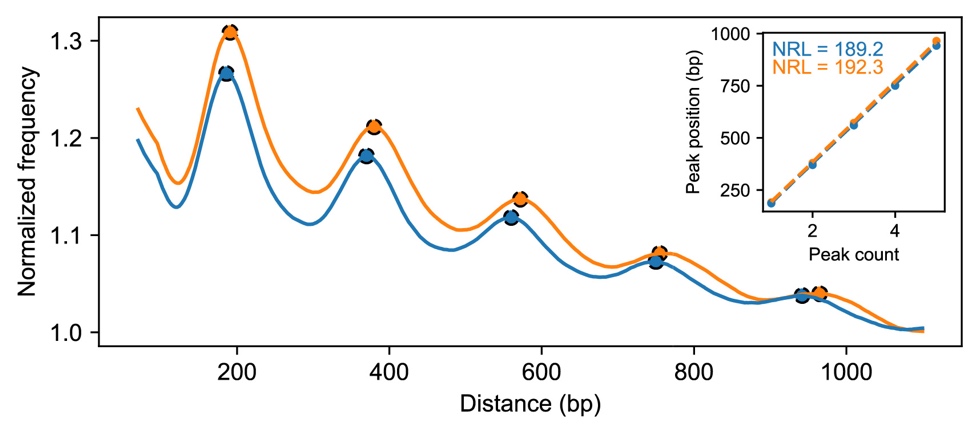

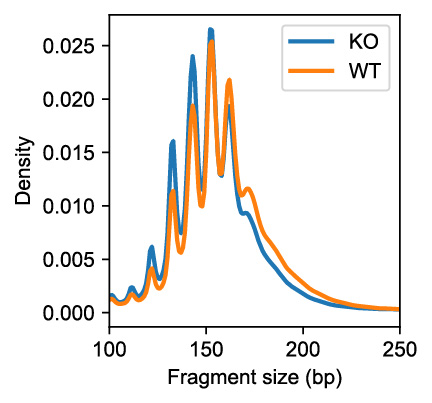

**Figure S7. Nucleosome spacing in euchromatin from H1.2 knockout and wild-type cells.**

**(Left)** Genome-wide mononucleosome-sized fragment (100-250 bp) distributions from MNase-seq of H1.2 knockout (KO, blue) and wild-type (WT, orange) HeLa cells. **(Middle)** Nucleosome positioning profile (phasogram) in euchromatin regions defined in blood cell types, as in Fig. 3, showing periodic spacing patterns. **(Right, inset)** Linear fit of nucleosome peak positions at euchromatin regions.

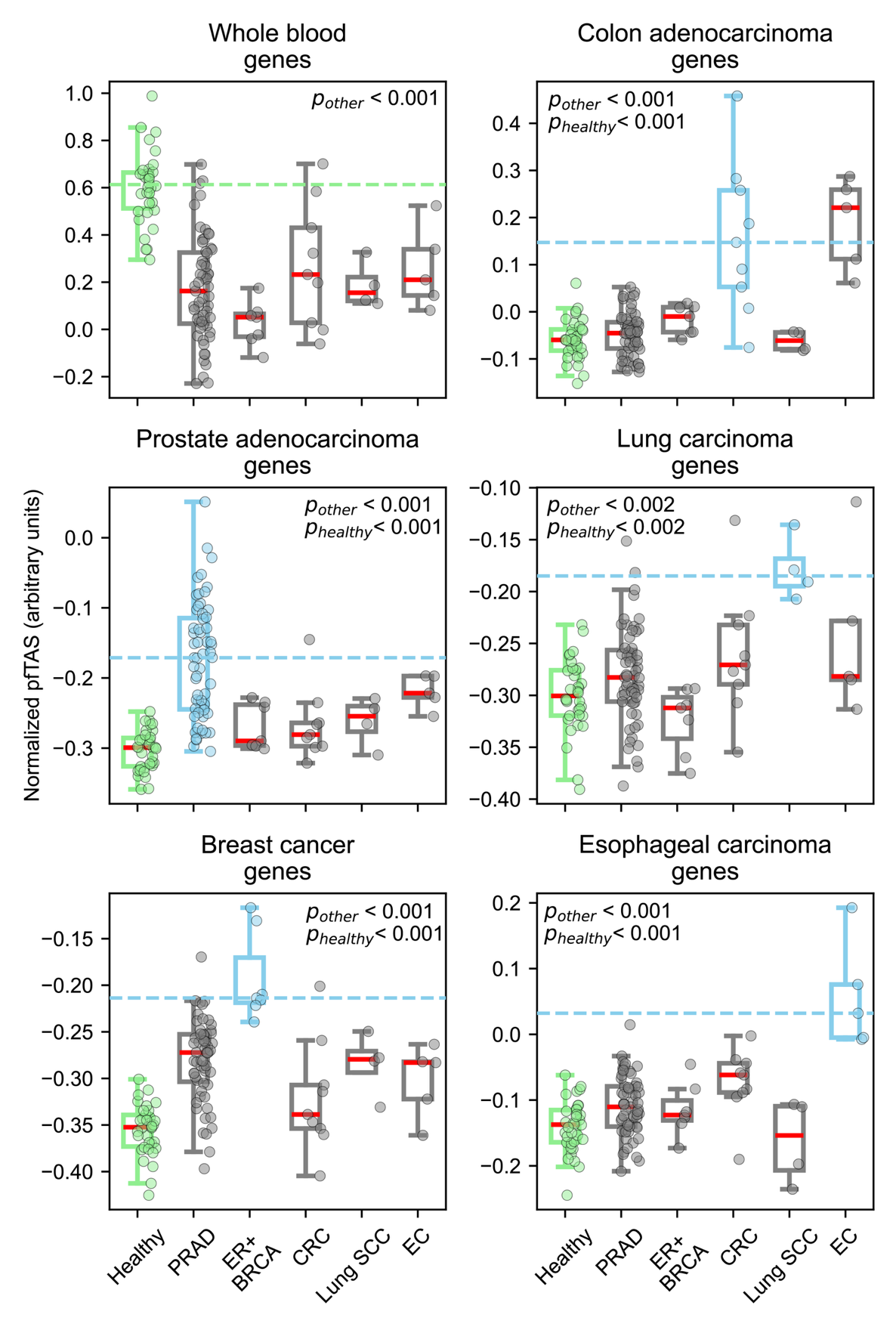

**Figure S8. cfDNA fragmentation patterns at lineage-associated genes.** A similar analysis is shown in **Fig. 4b**, but using the lineage-associated genes rather than their enhancers. pfTAS was calculated at each lineage-specific gene (from 8kb through the gene body). The boxplots represent median pfTAS for the indicated set of lineage-specific genes in cfDNA from patients with the indicated cancer. Light-green boxplots show values from healthy cfDNA for comparison. Boxplots colored in sky-blue indicate sample–gene set combinations where high expression is expected based on tissue of origin. Wilcoxon test p-values are indicated for comparisons of pfTAS between cancers in which high expression is expected versus pfTAS for other samples. *p_other_* indicates the *p*-value compared to other cancer samples, and *p_healthy_* indicates the *p*-value compared to healthy samples. The red solid line of each boxplot represents the median. Cancer types are abbreviated as follows: ER+ BRCA: ER-positive breast cancer, Lung SCC: squamous cell lung cancer, PRAD: prostate adenocarcinoma, CRC: colorectal cancer, EC: esophageal cancer.

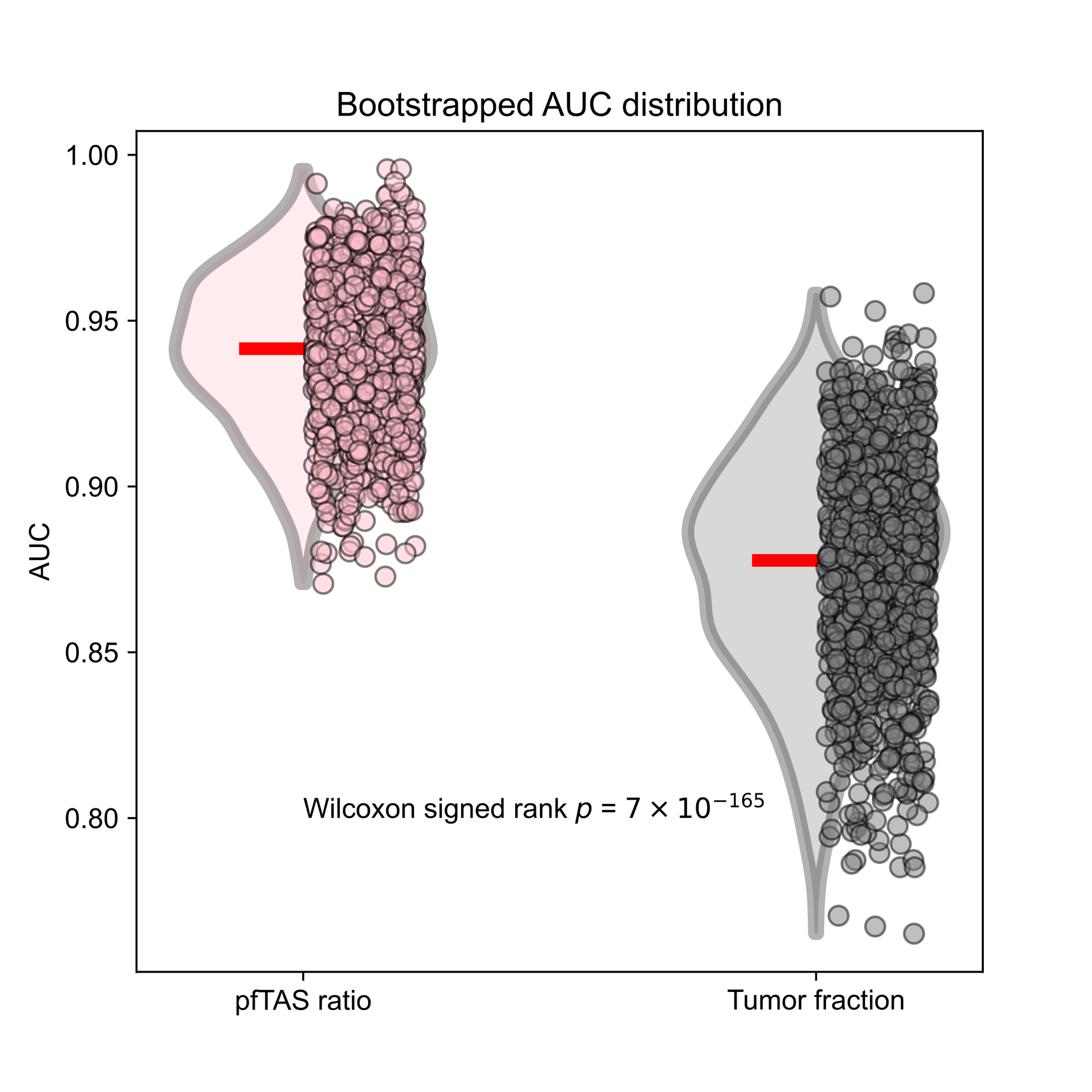

**Figure S9. Bootstrap distribution of AUC values for distinguishing SCLC and NSCLC**. The ratio of pfTAS SCLC regulatory elements vs NSCLC regulatory elements (see **Fig. 5c**), and cfDNA tumor fractions, were bootstrapped to calculate AUC distributions. Bootstrapping was repeated 1,000 times.

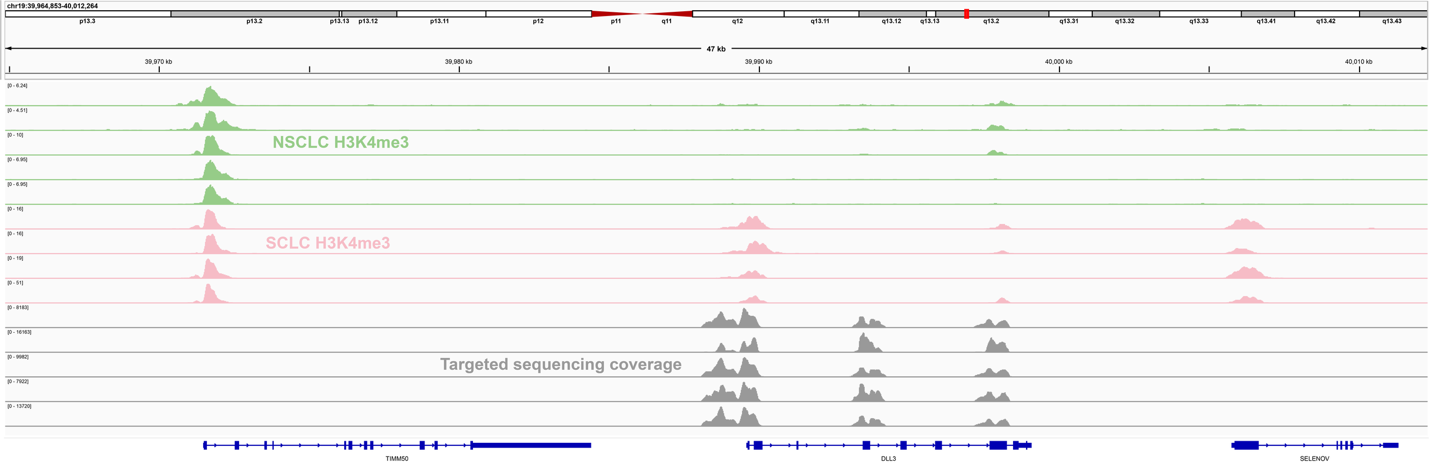

**Figure S10. IGV tracks of H3K4me3 ChIP-seq and targeted sequencing coverage**. The Integrative Genomics Viewer (IGV)^82^ was used to visualize normalized H3K4me3 coverage for NSCLC (light-green), SCLC (pink), and cfDNA fragments enriched by hybrid capture (gray) near *DLL3*. H3K4me3 coverage tracks for SCLC and NSCLC were retrieved from El Zarif et. al^36^. Published hybrid capture data from 5 samples^56^ were arbitrarily selected to examine coverage near *DLL3*.
